# FOXO3 regulated MIR503HG safeguards cellular quiescence by modulating PI3K/Akt pathway via miR-508/PTEN axis

**DOI:** 10.64898/2026.03.27.714688

**Authors:** Sonali R. Jathar, Vikas Dongardive, Juhi Srivastava, Vidisha Tripathi

## Abstract

Long noncoding RNAs (LncRNAs) have emerged as a class of important regulatory ncRNAs and are known to fine-tune numerous cellular processes including proliferation, differentiation and development; however, their role in quiescence still remains largely unexplored. A miRNA host gene lncRNA, MIR503HG, has been reported to play important role in cancer development. Here, we demonstrate the role of MIR503HG lncRNA in regulating cellular quiescence. MIR503HG displays elevated levels in human diploid fibroblasts induced to undergo quiescence. Depletion of MIR503HG in HDFs affects the entry of cells into quiescence but has no effect on cell cycle progression, suggesting its role in quiescence attainment and/or maintenance. Additionally, MIR503HG depletion led to a drastic decrease in the levels of miR508 target, PTEN with a concomitant increase in pAkt levels, indicating its role in negative regulation of miR508. Further, we demonstrate that the lncRNA MIR503HG regulates PTEN levels by acting as a ceRNA for miR508 to maintain cellular quiescence. Our studies illustrate that MIR503HG can function synergistically with miR503 to maintain cells under quiescence and both the miRNA-HG and the miRNA encoded by its gene locus synergistically control the same biological process in different ways by regulating different downstream genes.

## INTRODUCTION

The human body has about 10^13^-10^14^ cells; a greater part of which are in a non-dividing state. A few of these non-dividing cells are irreversibly arrested (senescent & terminally differentiated cells), while the rest of them have a capacity to re-enter the cell cycle depending upon physiological cues [1]. The latter group of reversibly arrested cells are termed as quiescent cells. Cellular quiescence is thus described as a reversible, non-dividing state prompted by varied anti-mitogenic cues, contact inhibition & various stresses [2]. This property is exhibited by various cell types, including adult stem cells, fibroblasts, lymphocytes, progenitor cells, hepatocytes, some epithelial cells & cancer stem cells[1–3]. On appropriate stimuli, cells may leave the quiescent state, re-enter the cell cycle and begin to proliferate. When stimulated, an intracellular signaling cascade drives global changes in gene expression accompanied by alterations in chromatin modifications that results in their rapid proliferation until required and then the cells exit the cell cycle and re-enter quiescence. This property of attaining quiescence of specific cells in multicellular organisms is essential for homeostasis, tissue regeneration and aging [4]. De-regulation of the balance between quiescence & proliferation can lead to various hypo- and hyper-proliferative pathologic conditions such as fibrosis, autoimmune diseases, neuro-degeneration, cancer and aging[5, 6]. A better understanding of this regulation has potential for identification of therapeutic targets. Recent studies have confirmed quiescence to be a highly active state characterized by distinctive transcription and epigenetic profile and metabolic status. For instance, various factors such as Rb, p53, p21, p27,p57, HES1, FOXO3 & many others have been discovered to play important roles in regulating quiescence[7–12]. Additionally, certain miRNAs such as miR-436, miR-126, miR-31 have been established to regulate the expression of specific genes in cells undergoing quiescence[13].

Another class of regulatory ncRNAs, long noncoding RNAs (lncRNAs) have been variously demonstrated to have a role in cell cycle regulation and cellular proliferation. It has been established that lncRNAs regulate various signaling pathways by modulating the function of key cell cycle regulatory genes and their dysregulated expression has been implicated in various disorders including cancer [14, 15]. However, their potential involvement in controlling cellular homeostasis or differentiation remains largely unknown. To understand the mechanisms by which lncRNAs govern the maintenance of a particular cellular state and how they are differentially regulated under various physiological conditions of a cell, a systematic analysis of lncRNA signatures in cells undergoing quiescence is key towards studying their function in the initiation and viability of the cellular state. Having a better understanding of functional roles of lncRNAs has tremendous potential to advance our understanding of cell regulatory and disease mechanisms. Various lncRNAs such as Linc-ROR, Gadd7, MEG-3, GAS5, Pnky have been shown to have an anti-proliferative function [16–18]. Also, MALAT1, HOTAIR, KCNQ1OT1, PANDA, SRA, lincRNA-p21 and many others are significantly involved in the cell cycle regulation either by controlling the expression levels of various cell cycle regulators or through direct interaction with important cell cycle regulators such as Rb & p53 [19]. Recently, PAPAS, antisense transcript from the rDNA promoter has been found to be up-regulated in quiescent cells. PAPAS interacts with Suv4-20h2 and guides it to nucleolar chromatin reinforcing quiescence mediated through H4K20me3 dependent chromatin compaction [20]. Overall, it can be stated that quiescence is a highly dynamic state which essentially requires the presence of certain regulators in the form of transcriptional regulators, epigenetic modifiers, post-transcriptional regulators to restrain the cell in a non-dividing state. We have recently demonstrated the role of a microRNA-host-gene lncRNA (lnc-MIRHG), MIR100HG, in cell cycle progression by modulating the activity of RNA binding protein HuR in an miRNA independent manner [21]. Similarly, MIR22HG functions as a tumor suppressor independently of miR-22 by acting as a ceRNA for miR-141-3p to augment DAPK1 levels in endometrial carcinoma [22]. Furthermore, several other studies have demonstrated miRNA independent roles of mature and processed lnc-MIRHGs. It has been reported that a significant fraction of miRNAs are encoded within lnc-MIRHGs in humans, which serve as diagnosis/prognosis markers [23–25]. Using an inducible classic model of human diploid lung fibroblasts for cellular quiescence, we performed a differential gene expression analysis of cells undergoing quiescence. Several lncRNAs were dynamically regulated during quiescence implicating the existence of an active gene regulatory network during quiescence. A recent study also reported a systematic transcriptomic profiling to identify differentially expressed protein coding and lncRNA genes during cellular quiescence and subsequent re-entry into the cell cycle in normal human diploid fibroblasts[26]. While this study further focussed on lncRNA genes that were differentially regulated during cell cycle re-entry, we compared the differentially expressed transcripts during quiescence with results from our laboratory. Among the differentially expressed lncRNAs in quiescent cells, MIR503HG displayed the most significant upregulation. MIR503HG has been variously reported to have an anti-proliferative function in several cancer cell lines [27, 28]. Moreover, the miRNAs encoded by this lnc-MIRHG, miR503 and miR424 have also been reported to negatively regulate key cell cycle regulatory genes [29–31]. A recent study demonstrated that MIR503HG was downregulated in non-small-cell-lung cancer and promoted cell apoptosis in vitro and repressed tumorigenesis in vivo[27]. Additionally, MIR503HG was upregulated in PE placental tissues compared to normal placental tissues and inhibited cell proliferation, invasion and migration of HTR-8/SVneo and JEG3 cells [32]. Furthermore, the involvement of MIR503HG in suppression of HCC metastasis by negative regulation of NFkB pathway was also reported recently [33]. MIR503HG has been demonstrated to have an anti-proliferative and anti-invasive function in bladder cancer [34]. In colon cancer, MIR503HG is downregulated and its overexpression inhibits proliferation, migration and invasion of colorectal cells mediated by TGF-b2 [35]. Increasing evidence demonstrates that the interaction between miRNAs and lncRNAs play a vital role in cancer. Various lncRNAs have been shown to share miRNA response elements (MREs) with certain mRNAs and act as competing endogenous RNAs (ceRNAs) to indirectly cross-regulate each other by competing for the same miRNAs. MIR503HG has been recently reported to inhibit TNBC tumor growth via the miR-224-5p/HOXA9 axis [28]. Additionally, in colon cancer cells MIR503HG inhibited the proliferation, migration and invasion via miR107/Par4 axis [36]. Taken together, all these studies have clearly established an anti-proliferative function of this RNA. However, the function of MIR503HG in normal cells or its involvement in cellular quiescence remains to be further explored. Moreover, significant upregulation of MIR503HG in quiescent normal human lung fibroblasts prompted us to investigate a mechanistic study to understand its involvement in quiescence regulation and largely cellular homeostasis.

We observed that the mature and processed MIR503HG acts independently and synergistically with miR503 to maintain cells in quiescent state. In our studies, we found that FOXO3 regulated MIR503HG is significantly up-regulated in quiescence condition. Further, functional studies revealed that MIR503HG acts independently of miR503 to maintain quiescent conditions by acting as a ceRNA for miR508 by regulating the Phosphatase and Tensin homologue (PTEN) levels. PTEN has been highlighted as an essential factor for safeguarding quiescence in adult muscle stem cells[37]. PTEN is a well-known tumor suppressor that dephosphorylates Akt, thereby suppressing the PI3K/Akt pathway. Constitutive activation of the PI3K/Akt pathway is one of the key features of various tumors that supports uncontrollable proliferation causing active tumorigenesis [38]. MIR503HG titrates the levels of miR508 thereby preventing PTEN degradation and finally facilitating cells into quiescence. Altogether, our studies recognize the novel role of the lncRNA MIR503HG in maintaining quiescence by regulating PI3K/Akt signaling pathway through miR508/PTEN axis.

### MIR503HG is upregulated in quiescent cells and promotes exit of cells from cell cycle into quiescent state

Cellular quiescence can be induced experimentally via serum starvation, loss of adhesion, or cell contact inhibition. We utilized normal human diploid lung fibroblasts WI38 cells to induce quiescence by serum starvation **[Fig 1A]**. WI38 cells become quiescent after 72-hour of serum starvation, reflected by the almost undetectable population of S-phase or G2/M phase cells **[Fig 1B-C]**. These cells can be reversed to enter cell cycle upon serum stimulation. Quantitative RT-PCR analysis of asynchronously growing and quiescent WI38 and NHDFs demonstrated a significant upregulation of MIR503HG in quiescent cells **[Fig 1D, S1A]**. We focused our investigation on MIR503HG for several reasons. MIR503HG was also detected in the quiescent population from a recently reported genome wide transcriptomic analysis on WI38 cells [26]. MIR503HG is transcribed from the X-chromosome and its gene locus encodes two miRNAs, miR503 and miR424, both of which were previously reported to have anti-proliferative roles. In addition, MIR503HG was also reported to have anti-proliferative function in some recent reports [27], [28], further implying that its upregulation during serum starvation could promote cellular quiescence. The differential expression of MIR503HG in quiescent cells implies that this RNA plays a vital role during this phase of the cell. Furthermore, we examined the levels of MIR503HG in various cell types and observed that it is upregulated upon induction of cellular quiescence in several cells **[Fig S1B]**. This strongly suggests that MIR503HG potentially is required for cellular quiescence. Biochemical fractionation of cells into nuclear and cytoplasmic components revealed that MIR503HG is predominantly nuclear both in asynchronously growing and quiescent cells **[Fig S1C-D]**. In order to determine the potential involvement of MIR503HG during cellular quiescence we analyzed the effect of its depletion in asynchronously growing WI38 cells. We employed ASO-mediated two rounds of knockdown within 24hrs interval and collected cells for PI-Flow cytometric analysis. Analysis of different sub-populations of cells in control and MIR503HG depleted cells revealed no significant change as the G1, S population of cells appeared comparable **[Fig 1F-G]**. PI-flow cytometry data does not segregate cells at G0 and G1 since both the subpopulations appear at the same location due to similar DNA content. To investigate if MIR503HG does have any effect on the quiescent cell population we stained cells with Hoechst-Pyronin and attempted to sort different sub-population of cells in control and MIR503HG depleted cells. However, due to the sensitive nature of WI38 cells and the requirement of bulk number of cells for sorting, we turned to HeLa cells for this experiment. As previously shown, MIR503HG is abundantly expressed in HeLa cells and gets upregulated when they are induced to undergo quiescence [**Fig S1B**]. In general, an asynchronously growing cell population consists of approx. 10% cells in quiescence. Interestingly, depletion of MIR503HG followed by Hoechst-Pyronin staining mediated cell sorting revealed reduction of the quiescent cell population with a concomitant increase in G1 population of cells **[Fig 1H]**. Further to confirm this, we synchronized WI38 cells to G0 by serum starvation in the presence and absence of MIR503HG and performed a flow-cytometric analysis. As expected, the control cells had efficiently undergone a quiescent state as revealed by almost negligible number of cells in S phase and G2/M phase, however, MIR503HG depleted cells were almost comparable to the asynchronous cell population even after serum starvation, further suggesting the requirement of MIR503HG for attainment of quiescence [**Fig S1E-F**]. Finally, we employed Hoechst-Pyronin based cell sorting in HeLa cells to identify the different cell populations upon serum starvation in control and MIR503HG depleted cells. As shown in **[Fig 1I]** while the control treated cells displayed majority of cells in G0 stage upon serum starvation, only approximately 10% cell population was in G0 stage upon MIR503HG depletion. Moreover, the G0 and G1 cell population of MIR503HG depleted samples upon serum starvation were almost comparable to asynchronously growing control, suggesting the involvement of MIR503HG in regulating quiescence. These observations were supported by reduced levels of the quiescence markers-HES1, p53 and MXI1 in MIR503HG depleted cells induced into quiescence **[Fig 1J]**. These results clearly confirm that MIR503HG has an important role during quiescence and is required for cells to exit from cell cycle into quiescence upon nutrient deprivation. To further determine if MIR503HG encoded miR503 is responsible for the exit of cells from cell cycle, we compared the levels of miR503 in quiescent cells and MIR503HG depleted cells. Depletion of MIR503HG resulted in almost 50% reduction in miR503 levels **[Fig. S1G]**. Additionally, we performed PI-flow cytometry analysis of the nutrient deprived (G0) cells treated with miR503 inhibitor. Although miR503 inhibitor treated cells did show some effect on the G0 subpopulation, however, it was not as comparable to MIR503HG depleted cells **[Fig S1H]**. This suggests that miR503 may not be solely responsible for driving the cells into quiescence and that MIR503HG does have an independent function in regulating the exit and entry of cells into quiescence.

**Fig. 1.**
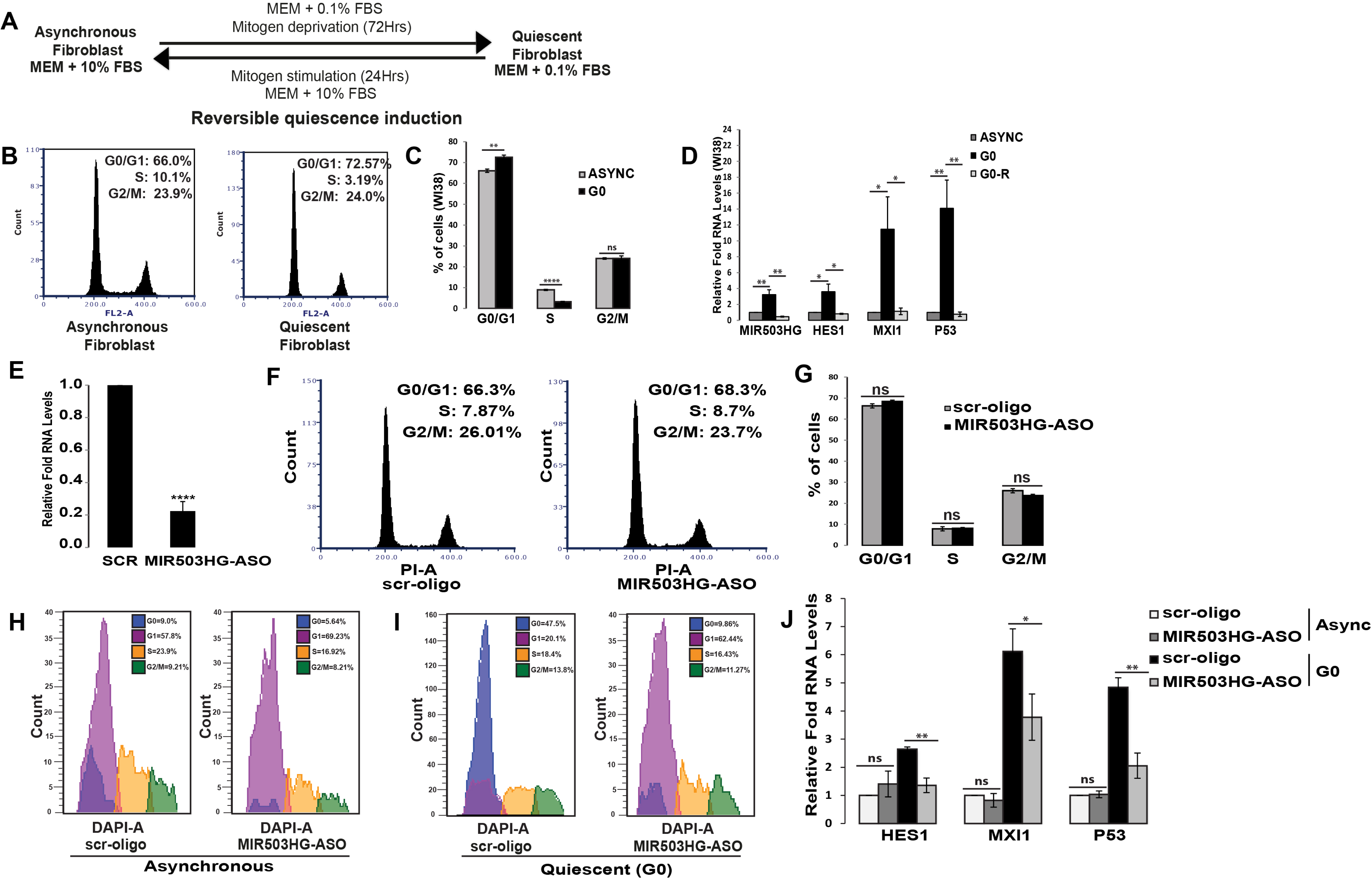
MIR503HG is upregulated in quiescent cells and promotes exit of cells from cell cycle into quiescent state: (A) Schematic representation of establishment of cellular quiescence induction model. (B) PI-Flow cytometry analysis of asynchronous and quiescent human diploid lung fibroblasts (WI38). (C) Quantitative representation of percentage cell population at each stage in asynchronous and quiescent WI38 cells. (D) RNA levels of MIR503HG, HES1, MXI1 and p53 examined by qRT-PCR in asynchronous, G0 and serum stimulated cells. (E) qRT-PCR analysis of MIR503HG RNA in control (scr) and MIR503HG depleted cells (MIR503HG-ASO) using antisense oligos. (F) PI-Flow cytometry analysis of WI38 cells treated with scr-oligo and MIR503HG-ASO. (G) Quantitative representation of percentage cell population at each stage in control and MIR503HG depleted cells. (H) Cell cycle profile of asynchronous HeLa cells treated with scr-oligo and MIR503HG-ASO distinguishing G0 and G1 populations using Hoechst-pyronin staining. (I) Cell cycle profile of serum starved quiescent HeLa cells treated with scr-oligo and MIR503HG-ASO distinguishing different cell populations using Hoechst-pyronin staining. (J) qRT-PCR analysis of quiescent markers HES1, MXI1 and p53 in asynchronous and quiescent cells treated with scr-oligo and MIR503HG-ASO. *: p≤0.05, **: p≤ 0.01, ***: p ≤0.001, ****: p ≤0.0001, ns: p> 0.05 by two-tailed student’s t-test, n=3. Error bars represent standard deviation.

### MIR503HG regulates key genes to maintain quiescent state

The lncRNAs are well known for regulating cellular processes by participating in numerous molecular pathways. For a systematic identification of pathways affected during quiescence entry and exit upon MIR503HG depletion, we performed a proteomic screen in asynchronously growing control and MIR503HG depleted cells, and also control and MIR503HG depleted cells that were induced to undergo quiescence. There was no significant overlap in the proteome profile between asynchronous and quiescent cells upon MIR503HG depletion. Comparison of the proteome data revealed a unique set of 319 proteins that were upregulated and 109 proteins that were downregulated upon MIR503HG depletion in quiescent cells **[Fig 2A]**. Majority of the upregulated proteins belonged to transcriptional and translational machinery (translation initiation, elongation, ribosome assembly, rRNA processing), indicating an active translation program upon MIR503HG depletion suggestive of pro-proliferative status of cells **[Fig 2B, Fig S2A]**. The quiescent cells display lowered metabolic activities, often characterized by decreased glycolysis. Upregulation of proteins required for various metabolic pathways including glycolysis, amino acid biosynthesis and fatty acid metabolism imply impairment of quiescence maintenance in MIR503HG depleted cells **[Fig S2B]**. Apart from this, proteins related to cell proliferation, MAPK pathway, mTOR pathway protein stabilization and export were also upregulated upon MIR503HG depletion **[Fig S2A, S2B]**. One of the major pathways that were affected in the down-regulated proteome upon MIR503HG depletion was the p38 MAPK pathway**[Fig S2C, S2D]**. p38 suppresses cell proliferation by a mechanism that involves increased expression of p21, which is CDK2 inhibitor [39]. Increased p21 levels in cells are an indicator of cell cycle arrest/cell quiescence [1], [3], [40]. Therefore, down-regulation of p38 pathways in MIR503HG depleted cells further suggests pro-proliferative status of the cells.

**Fig. 2.**
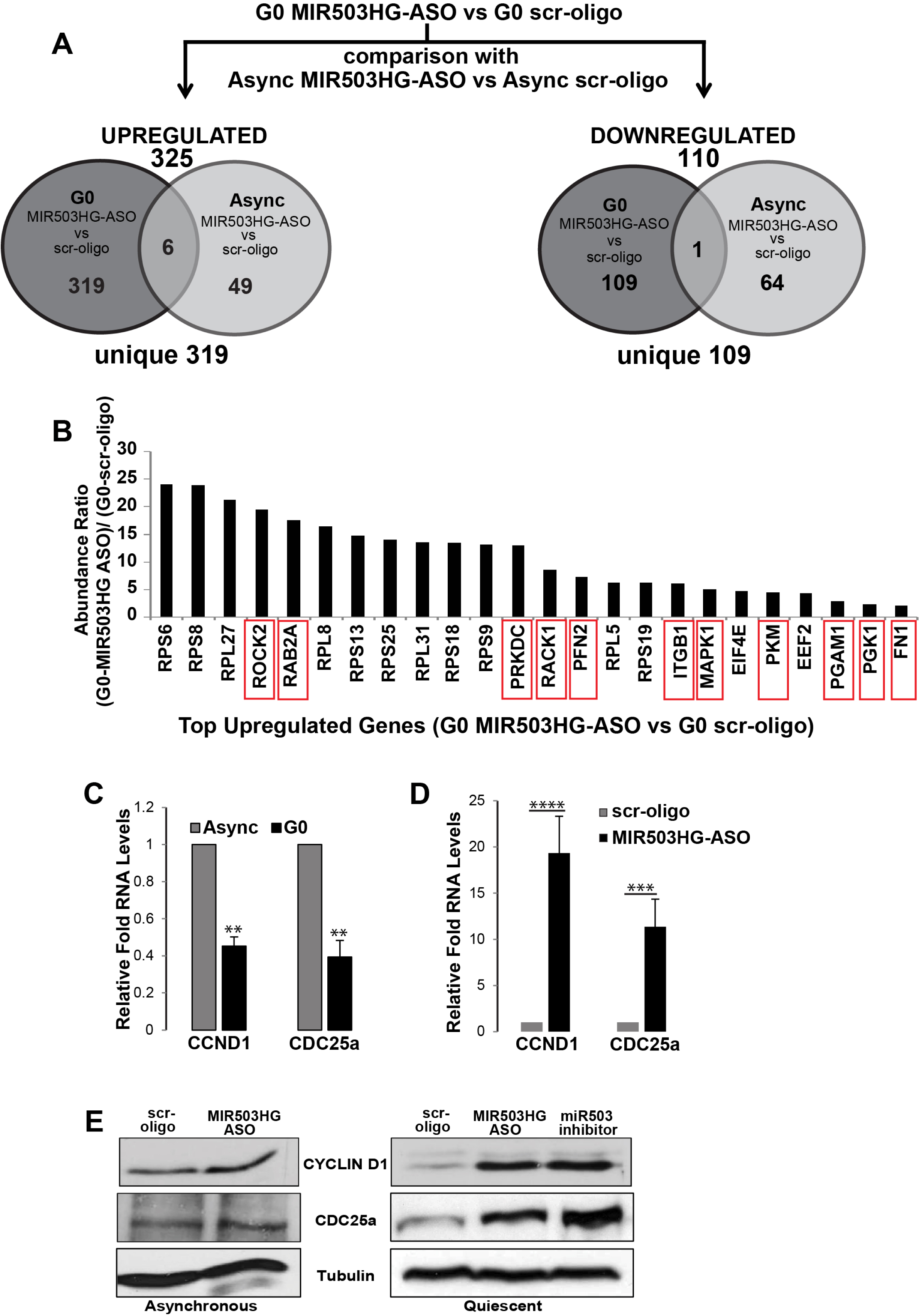
MIR503HG regulates key genes to maintain quiescence state: (A) Schematic representation of strategy employed for differential proteomic analysis in control and MIR503HG depleted cells in asynchronous and quiescent WI38 cells. (B) Top upregulated proteins in G0 cells upon MIR503HG depletion as compared to control. (C) qRT-PCR analysis of CCND1 and CDC25a RNA in asynchronous and quiescent WI38 cells. (D) qRT-PCR analysis of CCND1 and CDC25a RNA in control and MIR503HG depleted WI38 cells. (E) Cyclin D1 and CDC25a protein levels in control, MIR503HG depleted and miR503 inhibitor treated cells at asynchronous and quiescent conditions

It has been previously demonstrated that miR503 targets CCND1 and CDC25a to induce cell cycle arrest in cells [30, 41]. We also confirmed this by analyzing CCND1 and CDC25a both at RNA and protein levels in asynchronous and quiescent cells. Both CCND1 and CDC25a transcript levels were low in quiescent cells as compared to asynchronously growing WI38 cells **[Fig 2C]**; and as previously demonstrated, depletion of MIR503HG led to a significant increase in the transcript levels of both CCND1 and CDC25a **[Fig 2D]**. Western blot analysis of control and MIR503HG depleted cells revealed an increase in protein levels of CCND1 and CDC25a both in asynchronous and quiescent cell population, however, the increase in protein levels in quiescent cells was significantly more **[Fig. 2E]**. Additionally, quiescent cells treated with miR503 inhibitor displayed even further increase in the levels of both the proteins **[Fig 2E]**. Taken together, this data suggests that MIR503HG encoded miR503 targets CCND1 and CDC25a to maintain cells during quiescence. Finally, to confirm the overall quiescence phenotype regulated by MIR503HG encoded miR503, we transfected cells with miR503-mimic and performed a flow cytometric analysis. Interestingly, asynchronously growing cells transfected with miR503-mimic did show an increase in the G0/G1 population with a concomitant decrease in S phase cell population, however, the cells did not completely show a quiescent phenotype since there were approx. 10% cells still in S phase and 24% cells in G2/M **[Fig S2E]**. Altogether, this indicates that even though MIR503HG encoded miR503 mediates in regulating the levels of two key proteins CCND1 and CDC25a, MIR503HG has an independent role in the maintenance of quiescence phenotype of cells.

### MIR503HG is transcriptionally regulated by FOXO3/4

In order to understand the regulation of MIR503HG RNA during cellular quiescence, we performed genomic sequence analysis of MIR503HG and observed binding sites for transcription factors-FOXO3 and FOXO4 in its 5’ upstream regions. FOXO3 has been reported to serve as an essential transcription factor that promotes quiescence in adult stem cells by activating NOTCH signaling [45]. We speculated that FOXO3 transcriptionally regulates MIR503HG levels during quiescence. To confirm this, we transfected cells with FOXO3 and FOXO4 constructs and analyzed MIR503HG levels. In agreement with our assumption, we found that overexpression of both FOXO3 and FOXO4 leads to the up-regulation of MIR503HG which was comparable to increased levels of CDKN1B that is already known to be transcriptionally regulated by FOXO3 **[Fig 3A-B]**. Further we performed chromatin immunoprecipitation analysis to confirm binding of FOXO3 at MIR503HG promoter region in both asynchronously growing and quiescent cells. The immunoprecipitation data revealed enrichment of MIR503HG in FOXO3 immunoprecipitates, which was significantly more in quiescent cells **[Fig. 3C, S3A-B]**. This clearly indicates that FOXO3 binds to the MIR503HG promoter region and regulates its transcription. Higher enrichment of MIR503HG in quiescent cells suggests MIR503HG upregulation during quiescence mediated by FOXO3.

**Fig. 3.**
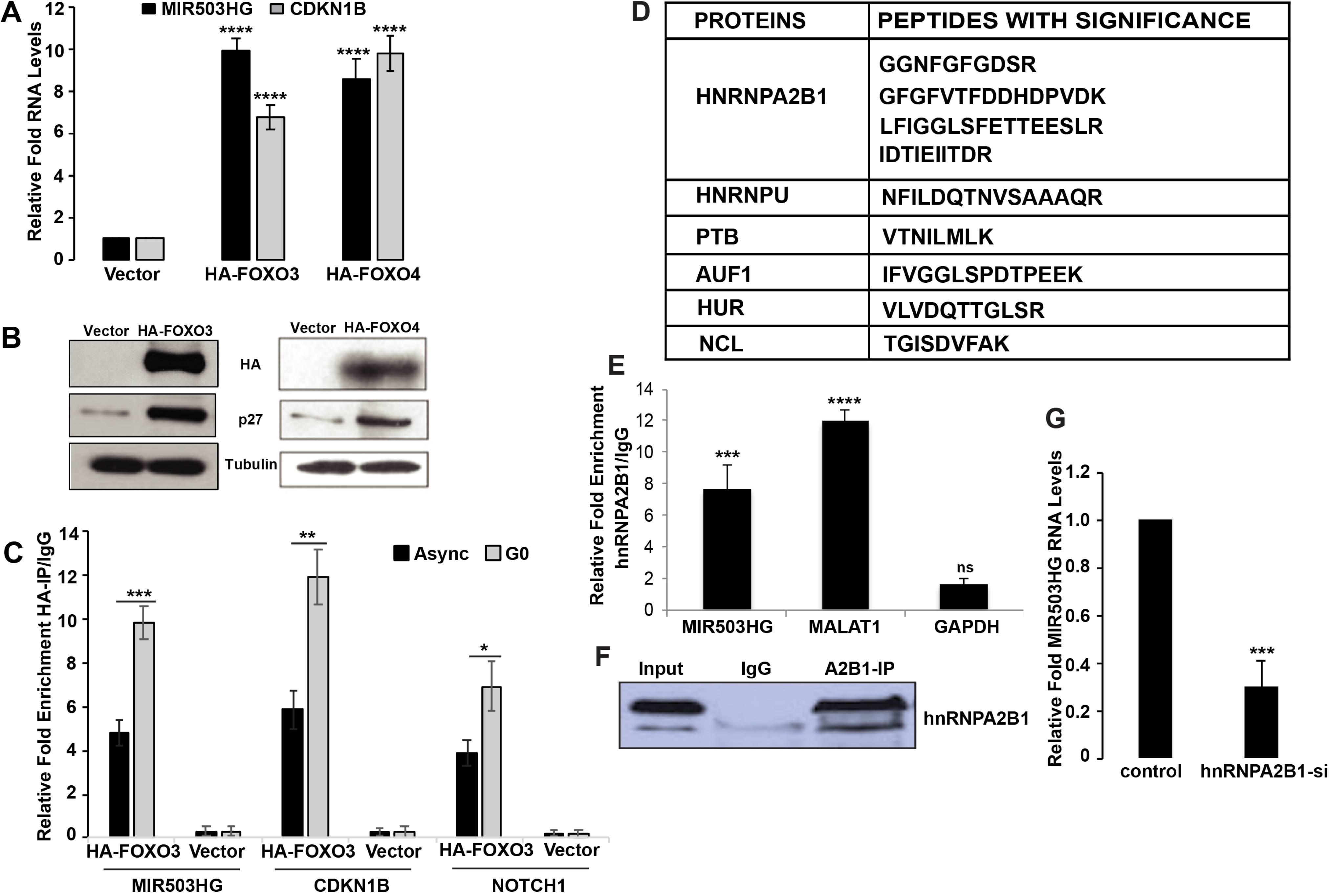
MIR503HG is transcriptionally regulated by FOXO3/4 and interacts with hnRNPA2B1: (A) qRT-PCR analysis for MIR503HG and CDKN1B in cells overexpressing vector control, HA-FOXO3 and HA-FOXO4. (B) Western blot analysis for FOXO3, FOXO4 and p27 in control and HA-FOXO3 or HA-FOXO4 transfected cells. (C) Chromatin immunoprecipitation using HA antibody to analyze relative fold enrichment of FOXO3 at MIR503HG, CDKN1B and NOTCH1 promoters in asynchronous and G0 cells. (D) List of proteins having peptides with significance upon mass spectrometric analysis of MIR503HG RNA pull down assay. (E) Relative fold enrichment of MIR503HG, MALAT1 and GAPDH upon RNA-IP using hnRNPA2B1 antibody. (F) Western blot analysis for hnRNPA2B1 in IP samples. (G) qRT-PCR analysis for MIR503HG RNA in control and hnRNPA2B1 depleted cells.

Several lncRNAs modulate protein function by interacting and forming an RNA-protein complex. In order to determine the molecular function of MIR503HG during cellular quiescence we investigated the MIR503HG-interacting proteins. We employed *in-vitro* transcribed biotinylated MIR503HG RNA and performed streptavidin pulldown in WI38 cell lysate followed by mass spectrometric analysis **[Fig. S3C].** From the protein interactome of MIR503HG, we identified hnRNPA2B1, HuR, PTBP proteins as top hits in our experiment [**Fig. 3D**]. Further, we validated the interaction of hnRNPA2B1 with MIR503HG by immunoblotting in the biotin RNA pulldown samples. hnRNPA2B1 RIP confirmed the interaction between endogenous protein and MIR503HG **[Fig 3E-F]**. To further investigate the functional relevance of the hnRNPA2B1 – MIR503HG interaction we analyzed the levels of hnRNPA2B1 in MIR503HG depleted cells and observed no significant difference in the levels of the protein. However, depletion of hnRNPA2B1 led to a significant reduction in the levels of MIR503HG RNA suggesting that hnRNPA2B1 may be required for the stability of this RNA**[Fig. 3G]**. Flow cytometric analysis of nutrient deprived hnRNPA2B1 depleted cells revealed subpopulations almost comparable to asynchronous cells further reiterating reduced MIR503HG levels in these cells **[Fig. S3D]**. Taken together, these results confirm that MIR503HG is transcriptionally regulated by FOXO3 during quiescence and is stabilized by hnRNPA2B1.

### MIR503HG titrates miR508 levels to promote cellular quiescence

LncRNAs normally act together with specific proteins or directly bind to miRNAs to regulate their function. Mounting evidence highlights the roles of numerous lncRNAs as competitive endogenous RNA (ceRNA) in modulating the expression or biological function of specific miRNAs in a wide variety of cellular contexts [43–45]. For instance, Linc-ROR has been described for its role in hESC self-renewal which is brought about by it acting as a ceRNA for the differentiation-related miRNAs such as miR-145 [43]. To further determine the parallel regulators of quiescence phenotype in cells induced by MIR503HG we turned to seek other miRNAs that may have an effect on cellular phenotype and possibly be regulated by MIR503HG. To identify MIR503HG miRNA targets of relevance to quiescence, we performed a computational analysis (miRanda & TargetScan) to screen for the miRNAs that share complementarity with MIR503HG **[Fig. S4A]**. Interestingly, we found complementary binding sites of miR191 and miR508 in the 3’ region of MIR503HG **[Fig. S4B, S4C]**. Most importantly, previous studies have demonstrated that miR508 targets PTEN, INPP5J and INPP4A [38]. miR508 directly suppresses these phosphatases resulting in constitutive activation of PI3K/Akt signaling [38]. Therefore, we hypothesized that MIR503HG might promote quiescence by sponging miR508 thereby upregulating PTEN. To test this, we depleted MIR503HG in asynchronous and quiescent WI38 cells. The alterations in PTEN, INPP5J and INPP4A RNA and protein levels were then measured with qRT-PCR and western blot analysis, respectively. As expected, knockdown of MIR503HG resulted in no significant change in PTEN and INPP5J levels in asynchronously growing cells **[Fig. 4A]**. However, transcript levels of PTEN and INPP5J were significantly high in quiescent cells as compared to asynchronous population. Most interestingly, depletion of MIR503HG in quiescent cells led to a significant reduction in the levels of PTEN and INPP5J **[Fig 4A]**. Moreover, this decrease in PTEN levels was also validated by western blot analysis of both PTEN and its downstream target protein AKT [**Fig. 4B**]. As indicated, MIR503HG depletion in quiescent cells displayed lower PTEN levels accompanied by an increase in phospho-AKT levels suggesting initiation of an active AKT signaling program in quiescent cells **[Fig. 4B]**. Moreover, treatment of miR508 inhibitor in asynchronous cell population did not result in any change in the pAKT levels, which further suggests that the negative regulation of miR508 via MIR503HG leading to increased PTEN levels is instrumental in maintaining quiescence. Taken together, these results confirm that miR508 is a downstream target of MIR503HG and driving factor of quiescence maintenance and implied a potential regulatory axis of cell fate. We further sought to investigate whether MIR503HG upregulated PTEN by binding and suppressing miR508 and thus inhibiting miR508 mediated PTEN suppression. To this end, miR508 inhibitor was transfected in control and MIR503HG depleted cells followed by qRT-PCR and immunoblot analysis for PTEN. As expected, the decrease in PTEN mRNA and protein levels induced by MIR503HG knockdown was rescued by miR508 inhibitor **[Fig 4C, S4D]** Consistent with this, miR508 inhibitor also resulted in elevated levels of proliferation markers and increase in cell proliferation as analyzed by flow cytometry in MIR503HG depleted cells **[Fig S4E]**. This suggests that MIR503HG induced cellular quiescence by inhibiting miR508. We further evaluated the effect of MIR503HG on miR508 levels and did not observe any significant difference in its levels between control and MIR503HG depleted cells **[Fig S4F]**.

**Fig. 4.**
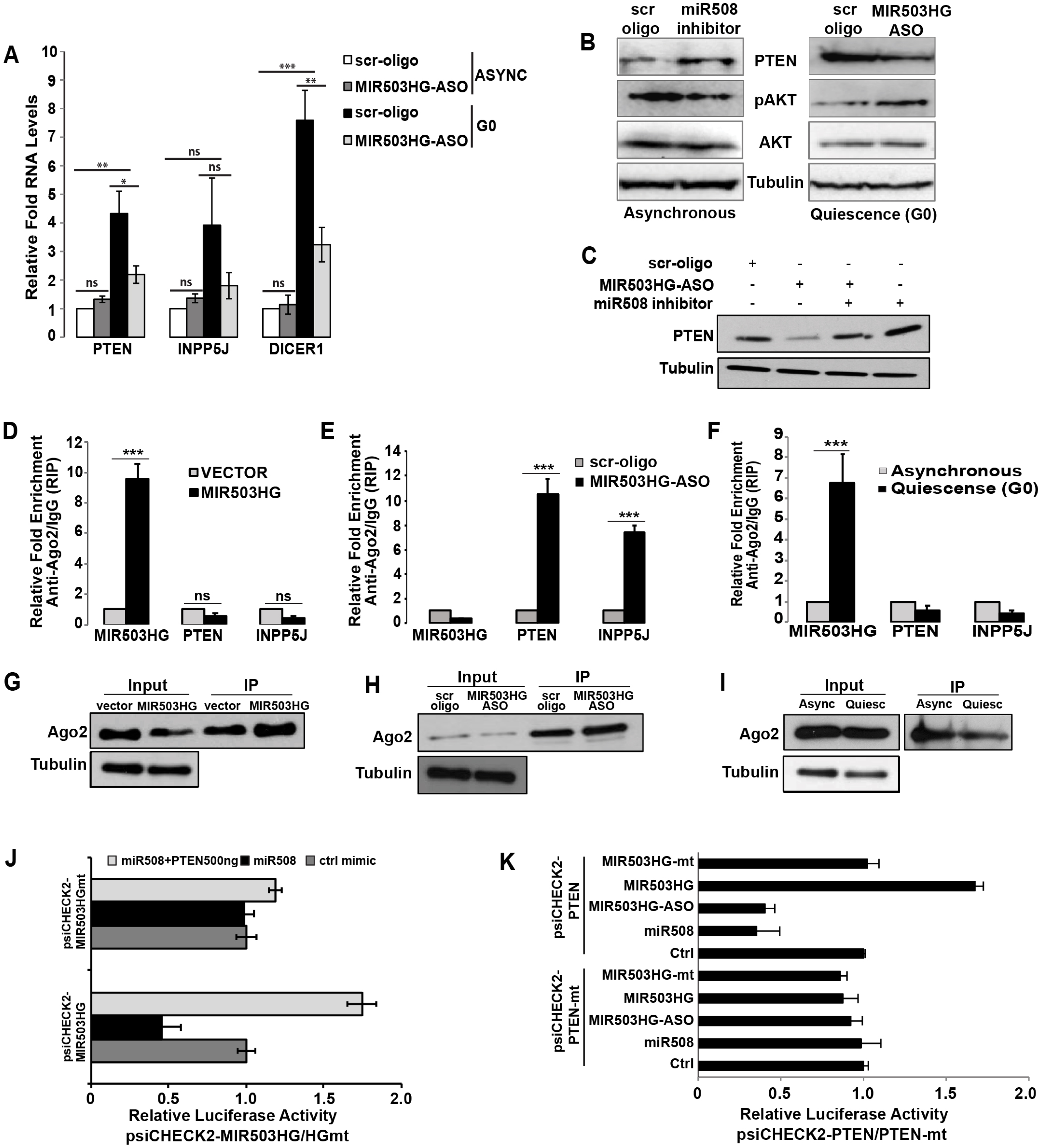
MIR503HG titrates miR508 levels to promote cellular quiescence: (A) qRT-PCR analysis for PTEN, INPP5J and DICER1 in asynchronous and G0 WI38 cells upon treatment with scr-oligo and MIR503HG-ASO. (B) PTEN, AKT and pAKT protein levels determined by immunoblotting in control WI38 and cells treated with miR508 inhibitor or MIR503-ASO. Tubulin was used as a loading control. (C) Western blot analysis to determine PTEN protein levels in WI38 cells treated with scr-oligo, MIR503HG-ASO or miR508 inhibitor. Tubulin was used as a loading control. (D) Ago2 RNA-immunoprecipitation assay in vector control and MIR503HG overexpressing cells followed by qRT-PCR analysis for MIR503HG, PTEN and INPP5J RNA. (E) Ago2 RNA-immunoprecipitation assay in control and MIR503HG depleted cells followed by qRT-PCR analysis for MIR503HG, PTEN and INPP5J RNA. (F) Ago2 RNA-immunoprecipitation assay in asynchronously growing and quiescent WI38 cells followed by qRT-PCR analysis for MIR503HG, PTEN and INPP5J RNA. (G) Ago2 RNA-immunoprecipitation assay in vector control and MIR503HG overexpressing cells followed by immunoblot analysis for Ago2. (H) Ago2 RNA-immunoprecipitation assay in control and MIR503HG depleted cells followed by immunoblot analysis for Ago2. (I) Ago2 RNA-immunoprecipitation assay in asynchronously growing and quiescent WI38 cells followed by immunoblot analysis for Ago2. (J) Relative luciferase reporter activity of psiCHECK2-MIR503HG and psiCHECK2-MIR503HGmt in cells transfected with miR508 mimic or co-transfected with miR508 mimic and PTEN. (K) Relative luciferase reporter activity of psiCHECK2-PTEN and psiCHECK2-PTENmt in cells transfected with miR508 mimic, MIR503HG-oe, MIR503HG-mt or MIR503HG-ASO.

LncRNAs can recruit and bind proteins to affect their cellular location, translational activity, or stability. Therefore, we analyzed the potential binding of MIR503HG and PTEN protein in our RNA-pull down assay. However, of all the enriched proteins identified in the assay, there was no significant enrichment of PTEN protein in the MIR503HG pull down sample compared to YFP-RNA control **[Fig. 3D]**. These results suggest that MIR503HG does not directly affect the stability of PTEN protein.

It is well established that miRNAs bind to their target transcripts to degrade mRNAs or inhibit protein translation in an Ago2-dependent manner. In order to determine whether MIR503HG was associated with RNA-induced silencing complex (RISC) containing miR508, we performed an Ago2 RNA-immunoprecipitation assay in control and cells overexpressing MIR503HG. As shown, MIR503HG was abundantly enriched in Ago2 pull down samples as compared to IgG control. PTEN and INPP5J did not show any significant enrichment upon Ago2 pull down **[Fig. 4D, 4G]**. Similarly, Ago2-IP in control and MIR503HG depleted cells displayed higher enrichment of PTEN and INPP5J upon pull down, implicating their association with RISC containing miR508 in the absence of MIR503HG **[Fig. 4E, 4H]**. Most importantly, Ago2-RIP assay in asynchronous and quiescent cells revealed higher enrichment of MIR503HG and lower enrichment of PTEN & INPP5J in quiescent cells as compared to asynchronous cells, similar to MIR503HG overexpression conditions **[Fig. 4F, 4I]**. Taken together, these results indicate association of MIR503HG with RISC containing miR508 and suggest that MIR503HG might function as ceRNA for miR508 in order to regulate PTEN and INPP5J levels during cellular quiescence.

To confirm a direct relationship between MIR503HG, miR508 and its other mRNA targets we performed luciferase reporter assay. Co-transfection of miR508 mimic and luciferase vector containing MIR503HG led to a significant decrease in the relative luciferase activity whereas not for mutant MIR503HG construct indicating that MIR503HG could directly modulate miR508. Additionally, ectopic expression of PTEN in the cells transfected with miR508 mimic diminished the negative effect of miR508 on the luciferase activity of wild type MIR503HG reporter**[Fig. 4J]**. Similarly, depletion of MIR503HG or co-transfection of miR508 mimic and PTEN luciferase reporter construct led to a significant decrease in the relative luciferase activity **[Fig. 4K]**. Moreover, overexpression of MIR503HG abolished the negative effect of miR508 on the luciferase activity of wild-type PTEN reporter **[Fig 4K]**. Overall, these results firmly reiterate that MIR503HG acts as ceRNA for miR508. It directly binds to miR508 thereby preventing its binding to the mRNA target PTEN and thus modulates PTEN levels in order to maintain cellular quiescence **[Fig. 5]**.

## Discussion

Quiescence is an important phenomena exhibited by various cell types, including adult stem cells, fibroblasts, lymphocytes, progenitor cells, hepatocytes, some epithelial cells & cancer stem cells [1, 2]. The balance between quiescence and proliferation should be maintained for a proper tissue homeostasis [1, 4]. Any disturbance in this balance leads to various hyper- and hypo-proliferative diseases [1]. Proliferation has been widely studied and the factors involved in proliferation regulation are well known. Whereas, extensive research needs to be done in order to delineate quiescence regulation. LncRNAs have emerged as a class of important regulatory ncRNAs and are known to fine-tune numerous cellular processes such as cell cycle regulation, splicing, dosage compensation, development etc [15, 19, 46, 47]. Several lncRNAs such as MEG3, GAS5 have been reported in anti-proliferative roles [17, 48]. We aimed to understand the involvement of lncRNAs in quiescence regulation. A detailed expression study from our laboratory revealed a unique set of quiescence specific lncRNAs showing higher fold expression levels in quiescent fibroblasts as compared to asynchronous fibroblasts. One of the transcripts that displayed a dramatically high level in quiescent cells was MIR503HG.

MIR503HG is located on chromosome Xq26.3 and is a host gene for miR503 and miR468. We have previously demonstrated miRNA-independent roles of mature and processed lnc-MIRHGs in cellular proliferation and cell cycle regulation. Moreover, studies from several other groups strongly suggest lnc-MIRHGs can serve as diagnosis/prognosis markers and play regulatory roles in various cellular processes. Recently we have demonstrated that MIR100HG regulates cell cycle progression by modulating the activity of RNA binding protein HuR in a miRNA-independent manner. Similarly, a recent study reported a systematic transcriptomic profiling of differentially expressed protein coding and lncRNA genes during cellular quiescence and subsequent re-entry into the cell cycle in normal human diploid fibroblasts. Interestingly, several dynamically regulated MIRHGs were expressed in quiescent cells. In the current study, we demonstrate that MIR503HG is upregulated in quiescent cells, and the mature and processed MIR503HG acts independently and synergistically with miR503 to maintain cells in quiescence state.

Several recent reports have demonstrated an anti-proliferative role for lncMIR503HG and established the function of miR503 as a negative regulator of cell proliferation. Upregulation of both these transcripts in quiescence cells is indicative of their potential role in regulation of this program. We confirmed this by transient depletion of MIR503HG in cells induced to undergo quiescence. The fact that there was no effect on the cells already committed to the cell cycle but a significant reduction in the quiescent population suggested that MIR503HG is required for maintenance of cells in quiescence. Depletion of MIR503HG using phosphorothioate modified antisense-oligonucleotides resulted in more than 80% knockdown of the lncRNA and approx. 50% reduction in miR503 levels. However, there was no significant effect on normal cell cycle progression upon knockdown. This indicates that the reduced levels of miR503 achieved were not enough to lead any significant effect on cell cycle progression.

The FOXO family (FOXO1, FOXO3, FOXO4 and FOXO6) of transcription factors are known to regulate varied cellular programmes, including cell survival, proliferation, differentiation as well as metabolism [49]. Moreover, FOXO3 transcription factor shows significant expression in quiescent muscle stem cells and promotes the stem cell quiescence by activating Notch signaling during adult muscle regeneration [39]. Additionally, FOXO3 maintains the adult neural stem cell (NSC) pool by triggering a gene cascade that ensures preservation of quiescence [8]. Based on the ChIP data, we observed higher enrichment of FOXO3 at MIR503HG promoter in quiescent cells suggesting that upregulated levels of MIR503HG during cellular quiescence is a result of transcriptional activation mediated through FOXO3 transcription factor. Heterogeneous nuclear ribonucleoprotein A2B1 (hnRNPA2B1) belongs to hnRNP family of proteins and has been reported to bind to RNAs and regulate their transcription, stability, splicing and transport from nucleus to cytoplasm [50]. Our studies confirmed that hnRNPA2B1 binds to and stabilizes MIR503HG. It is established that the PI3-kinase/Akt signaling promotes cell survival by inhibiting p38 mitogen-activated protein kinase (MAPK)-dependent apoptosis. Blockage of PI 3-kinase or Akt pathways by serum withdrawal in cells stimulates p38-dependent apoptosis. The molecular mechanisms underlying cellular quiescence programs indicate that activation of multiple signaling pathways, chromatin remodelling and other epigenetic factors drive a cell to undergo a reversible state of quiescence upon environmental cues. The p38 pathway is one of the major mitogen-activated protein kinase pathways and is a mediator of inflammation, stress response, tumor suppression and oncogene induced senescence. P38 suppresses cell proliferation by a mechanism that involves increased expression of CDK2 inhibitor, p21. An increase in the levels of p21 is indicative of cell cycle arrest and quiescence. Interestingly, proteome analysis of the cells depleted of MIR503HG revealed downregulation of the p38 pathway in cells induced to undergo quiescence, suggesting a pre-proliferative status of the cells. A detailed analysis of the protein cohort in MIR503HG depleted quiescent cells revealed upregulation of proteins related with cell proliferation, MAPK pathway, mTOR pathway, protein stabilization and export. This data further reiterates that despite the environmental cue of serum withdrawal, depletion of MIR503HG itself could change the gene expression program towards cell proliferation.

Mounting evidence suggests that lncRNAs can associate with proteins or other RNAs, for instance, miRNAs and function as miRNA sponges to further affect the expression of miRNA target genes. For example, the lncRNA NORAD have been demonstrated to regulate lung cancer cell proliferation, apoptosis, migration and invasion by the miR30a/ADAM19 axis. Using bioinformatic prediction tools we identified several potential miRNA targets for lncMIR503HG i.e. miR191, miR508, miR4765, miR1237 etc. Interestingly, the known mRNA targets of miR508 are PTEN, INPP4A and INPP5J, which are protein phosphatases that negatively regulate the PI3-Akt signaling pathway. PTEN is a tumor suppressor and an essential regulator of cellular quiescence. Based on this, we sought to understand if MIR503HG regulates PTEN levels in cells to maintain quiescence by titrating/sponging miR508 levels. The luciferase assay and Ago2-IP confirmed our hypothesis. Additionally, MIR503HG depletion in quiescent cells displayed significantly reduced levels of PTEN with a concomitant increase in phospho-Akt levels indicating exit of cells from quiescence into proliferation. Activation of Akt signaling also indicates the inhibition of p38-MAPK pathway that we observed in our proteomic analysis upon MIR503HG depletion. In conclusion, MIR503HG which is often downregulated and co-expressed with miR503, is upregulated by FOXO3 mediated transcriptional activation during quiescence and its higher levels sequester the miR508 pool thereby restricting PTEN inhibition. The stabilization of PTEN leads to establishment and maintenance of the quiescence program. Moreover, MIR503HG can function synergistically with miR503 to maintain cells under quiescence. Both the lnc-MIRHG and the miRNA encoded by its gene locus synergistically control the same biological process in different ways by regulating different downstream genes.

## ACKNOWLEDGEMENT

We would like to thank V. Tripathi lab members for their comments and suggestions. We thank Dr. A Chakraborty for his help in mass spectrometric analyses, luciferase assays and ChIP assays. We thank NCCS FACS facility for Flow cytometry experiments. This work was supported by Early Career Research Grant from Science and Engineering Research Board-DST (SERB-ECR; ECR/2015/000242); DBT-Ramalingaswami re-entry fellowship (BT/RLF/Re-entry/01/2013) and NCCS intramural funding.

## MATERIALS AND METHODS

### Cell lines and Reagents

The cell lines-WI-38, HeLa, U2-0S, MCF7, and HEK293 were purchased from ATCC. The NHDF cell line was purchased from Lonza. All the reagents were of analytical grade and were purchased from Invitrogen, Sigma and Hi-media.

### Mammalian Cell Culture

The normal human diploid lung fibroblast, WI-38 (American Type Culture Collection, Cat no-ATCC-CCL-75) was cultured in Minimum Essential Medium (MEM)(GIBCO) supplemented with 10% FBS (GIBCO), 0.1% Non-essential amino acids (MP Biomedicals), 0.1% Penicillin-Streptomycin (GIBCO) at 37°C with 95% relative humidity and 5% CO2. The HeLa cells were cultured in high glucose Delbucco’s Modified Eagle’s Medium (DMEM)(GIBCO) supplemented with 10% FBS (GIBCO), 0.1% Penicillin-Streptomycin (GIBCO) at 37°C with 95% relative humidity and 5% CO2. For sub-culturing, cells were trypsinized by 0.25% Trypsin-EDTA [for WI38 cells] and 0.05 Trypsin-EDTA [for HeLa] respectively. The trypsin was neutralized by adding complete medium and the cells were subsequently pelleted down at 2000xg for 3 minutes. Cell pellet was then re-suspended in fresh culture medium and plated in cell culture dish as per the desired confluency.

### Quiescence Induction

The asynchronously growing WI-38 and HeLa cells in high glucose medium (MEM/DMEM) + 10%FBS were washed with PBS (cell culture) and then incubated in medium (MEM/DMEM) containing 0.1% FBS for 48 hours. The cell population was analyzed through Propidium Iodide-Flow Cytometry and Hoechst-Pyronin flow cytometry for proper induction of quiescence.

### Subcellular Fractionation

The subcellular fractionation was performed by biochemical method. The asynchronous and quiescent cells were first washed with ice cold 1X PBS twice and then pelleted down at 1200 rpm for 5 minutes. The pellet was resuspended in 1ml of ice-cold lysis buffer B (10Mm Tris (pH=8.4), 140 mM NaCl. 1.5 mM MgCl2, 0.5% NP40, 1mM DTT, 200U/ml RNase inhibitor) with slow pipetting (usually around 50 ups and down using 1 ml pipette) with no visible clumps followed by centrifugation at 1000xg at 4 °C for 5 minutes. The resultant supernatant (cytoplasmic fraction) was resuspended in 1 ml Trizol LS. The pellet was again resuspended in 1ml lysis buffer B along with 1/10th volume (100ul) of detergent stock (3.3% (w/v) sodium deoxycholate, 6.6% (v/v) Tween20) under slow vortexing followed by incubation on ice for 5 minutes. After the incubation, the resuspension was centrifuged at 1000xg at 4 °C for 5 minutes. The resultant supernatant (postnuclear fraction-ER/cytoplasmic fragments) was added to the previously collected cytoplasmic fraction. The pellet (nuclear fraction) was rinsed with lysis buffer B once and then re-suspended in Trizol.

### MIR503HG Knockdown

Phosphorothioate internucleosidic linkage-modified antisense oligonucleotides [ASOs] were used to deplete human MIR503HG. The oligonucleotides were transfected to cells two times (48hr) within a gap of 24hrs, at a final concentration of 50nM, using Lipofectamine RNAimax reagent as per the manufacturer’s instructions (Invitrogen, USA). The knockdown efficiency was analysed through qRT-PCR and the effect on cell proliferation was studied by PI-Flow Cytometry.

### Real Time PCR

Total cellular RNA was extracted from cells using Trizol (Invitrogen) as per manufacturer’s instructions and reverse transcribed into cDNA using Multiscribe Reverse transcriptase and Random Hexamers (Applied Biosystems). qPCR was performed using StepOne Plus system (Applied Biosystems) and Quant Studio Flex 6. Transcript levels were quantitated against a standard curve by Real-time RT-PCR using the SYBR Green I fluorogenic dye and data analyzed using the StepOne plus system software. Primer sets showing comparable high efficiencies were used for the analyses.

### Propidium Iodide (PI) Flow Cytometry

Cells were collected and washed in chilled PBS, re-suspended in 1X PBS and fixed in ethanol and stored at 4°C for minimum of 1 hour or until further processing. Later, the fixed cells were washed with 1X PBS and re-suspended in PBS with 10µg/ml RNaseA, 0.1% TritonX-100 and 20µg/ml Propidium Iodide (PI), and further incubated at 37°C for 30 minutes. The DNA content was measured by flow cytometry on FACS Calibur (BD Biosciences) using FL2 filter and analysis was performed with the help of Cell Quest Pro software.

### Hoechst-Pyronin-Y Flow Cytometry

The HeLa cells were seeded at ∼40% confluency for two rounds of knockdown. After, two rounds of knockdown, the cells were collected with and without quiescence induction. For Hoechst/Pyronin FACS, the live cells were first re-suspended in pre-warmed culture medium containing Hoechst dye (10ug/ml) and further incubated at 37°C for 45 minutes. Later, Pyronin Y (0.5ug/ml) was added to the same cell suspension, followed by 45 mins incubation at 37°C. The DNA and RNA content of the cells was measured on FACS ARIA III SORP (BD Biosciences) and the analysis was performed with the help of Diva software.

### SDS-PAGE and Western blotting

Cells were scraped in medium and washed with ice-cold PBS. Extraction was performed in lysis buffer (80mM Tris-Cl (pH=6.8), 15% glycerol, 2%SDS, 100mM DTT) containing protease and phosphatase inhibitors for 15min on ice. Loading dye (250mM Tris-Cl (pH=6.8), 45% glycerol, 5%SDS, 0.01% Bromophenol blue, 5% beta mercaptoethanol) was directly added to the lysate and boiled for 5min and finally loaded onto the polyacrylamide gel. The SDS gel was run using electrophoresis gel running buffer (25Mm Tris, 0.1% SDS, 250Mm glycine, pH=8.3) at 120V. Western blotting was performed as described previously (Tripathi et al., 2012). The resolved proteins were transferred on to the nitrocellulose membrane (Merck Millipore) using wet transfer unit (GE Biosciences) with carbonate transfer buffer (250mM NaHCO3, 170mM Na2CO3, 20% methanol, pH=9.5) for 3 hours. After the transfer, the protein transfer was confirmed by staining the membrane by 0.1% Ponceau staining solution. The membrane was then blocked using 5% skimmed milk made in TBST (100Mm Tris-Base, 0.9% NaCl, 0.1% Tween20) or 3% BSA made in TBST for 1 hour. The membrane was then incubated with the primary antibody [Cyclin D1(1:100), CDC25a(1:100), p27(1:1000), Ago2(1:1000), PTEN(1:500), Akt(1:500) t, p-Akt(1:500), Tub (1:5000)] prepared in 5% skimmed milk or 3% BSA in TBST at 4 °C O/N. After the incubation with primary antibody, the membrane was washed with TBST thrice for 10 minutes each. The membrane was then incubated with secondary antibody [anti-mouse HRP conjugate (1:5000) from Cloud clone] for one hour at room temperature, followed by three TBST washes 10 minutes each. The proteins were detected using ECL chemistry on GE Amersham.

### Biotin RNA pulldown assay

Biotin labeled RNA was synthesized as per the manufacturer’s instructions (Biotin RNA Labeling Mix, Roche). MIR503HG full-length cDNA clone in pCR2.1-TOPO (Invitogen, USA) vector were linearized as DNA template during in vitro transcription. In each reaction, 1ug linearized plasmid was used, and T7 RNA polymerase was used for biotin-labeled RNA synthesis. In vitro transcription was done at 30°C for overnight followed with DNase I (Sigma) treatment as per the manufacturer’s instruction. For the detection of interacting proteins, 107 WI38 cells were collected in pellet and resuspended in 2mL PBS, 2 mL nuclear isolation buffer (1.28M sucrose, 40mM Tris-HCl pH=7.5, 20mM MgCl2, 4%Triton X-100) and 6mL H2O on ice for 20min with frequent mixing. Nuclei were pelleted by centrifugation at 2500g for 15min at 4°C. Nuclear pellet was resuspended in 1mL RIP buffer (150mM KCl, 25mM Tris pH=7.4, 0.5mM DTT, 0.5%NP40) containing protease inhibitors, and incubated on ice for 10min. Cells were further lysed using a Dounce homogenizer with 15-20 strokes. Debris was then removed by centrifugation at 13000rpm for 10min. For pulldown, 40uL Dynabeads M-280 Strepatavidin was washed 2 times with Buffer A (0.1M NaOH, 0.05M NaCl), and with Buffer B (0.1M NaCl). 1mL of nuclear extract was added to beads together with 1ug biotin-labeled folded RNA probe, RNase Inhibitor (Invitrogen), 5 uL 50 mg/mL Heparin (Sigma, H3149), and 5uL Yeast t-RNA (Sigma, R5636) for pre-clearing at 4°C for 1hr. Pre-cleared nuclear extract was then added to pre-washed beads for pulldown at 4°C for 1.5hr. Beads then washed with high salt buffer (0.1%SDS, 1%Triton X-100, 2mM EDTA, 20mM Tris-HCl pH=8.0, 500mM NaCl), low salt buffer (0.1%SDS, 1%Triton X-100, 2mM EDTA, 20mM Tris-HCl pH=8.0, 150mM NaCl) and TE buffer (1mM EDTA, 10mM Tris-HCl pH=8.0). 40uL protein loading buffer was then added to the beads and heated at 95°C for 10min for protein samples. The protein samples were electrophoresed on 10% SDS-PAGE, silver stained as per manufacturer’s instructions (Invitrogen) and further subjected to mass spectrometric analysis (MS). The MS data was further confirmed by immunoblotting using specific antibodies.

### In-silico miRNA binding site prediction

In order to identify the miRNAs targeting MIR503HG, the binding of miRNAs (miRBase) to MIR503HG was predicted using two bioinformatics prediction tools-TargetScan and miRanda, keeping the free energy cutoff <-20. The putative miRNA binding sites discovered with the help of both the prediction tools were compared with each other and only the common predictions were considered for the subsequent experiments. The discovered miRNA binding sites were also confirmed by curating StarBase v2.0 database Li et al., Nucleic Acids Research, January 2014.

### Luciferase assay

The wild-type and mutant MIR503HG and PTEN were cloned in psiCHEK2 vector. For luciferase reporter assay, firefly luciferase vector and the respective wild type (psiCHEK2-MIR503HG, psiCHEK2-PTEN) or mutant psiCHEK2 (psiCHEK2-MIR503HGmt, psiCHEK2-PTENmt) constructs were co-transfected in HeLa with miR-508 mimic or negative control using Lipofectamine 2000 (Invitrogen). The relative firefly luciferase activity was measured after 48 hours of transfections and was normalized to Renilla luciferase activity (Promega).

### Ago2 IP

Ago2 immunoprecipitation was performed for MIR503HG overexpression, MIR503HG knockdown, asynchronous, and quiescent conditions in HeLa cells. Briefly, after MIR503HG OE, MIR503HG knockdown, and quiescence induction, the cells were lysed (Tris-HCl pH=8 50mM, Nacl 150 mM, NP-40 1%, Sodium deoxycholate 0.5%, SDS 0.1%) and centrifuged at 12,000xg for 30 minutes at 4°C. Anti-ago2 antibody was then added to the lysate at the dilution of 1:150 (1µl antibody per 150µl lysate) and the lysate was incubated O/N on rotation at 4°C. After the O/N incubation, 80µl protein G agarose beads (Pierce™ Protein G Agarose) were added to the lysate, followed by 2 hours incubation on rotation at 4°C. The beads were then washed thrice with 1X PBS on rotation at 4°C for 10 minutes each. The washed beads were then processed for RNA isolation and qRT-PCR for MIR503HG, PTEN and INPP5J.

### Chromatin Immuno-precipitation (ChIP)

ChIP was performed for assessing the binding of FOXO3 to promoter region of MIR503HG in both asynchronous and quiescent conditions with HA-FOXO3 overexpression. Briefly, the cells were first crosslinked with formaldehyde (final conc.∼0.75%) with gentle rotation at RT, followed by addition of glycine (final conc.∼125mM) and incubation at RT for 5 minutes to quench formaldehyde. The cells were then washed with ice cold 1X PBS, followed by scraping and centrifugation at 1000xg for 5 minutes at 4°C. The cells were then re-suspended in ChIP lysis buffer (50 mM HEPES-KOH pH7.5, 140 mM NaCl, 1 mM EDTA pH8, 1% Triton X-100, 0.1% Sodium Deoxycholate, 0.1% SDS, protease inhibitor) and incubated on ice for 10 minutes. The lysed cells were sonicated to get 200-1000bp size fragmented DNA and centrifuged at 8000xg for 10 minutes at 4°C to pellet down the cell debris, while the supernatant (chromatin preparation) was used for ChIP. Anti-HA antibody (5 µg per 20 µg DNA) was added to the chromatin preparation and incubated on rotation at 4°C for 2 hours. Then 50 µl of protein G beads (Pierce™ Protein G Agarose) were added to the samples and incubated on rotation at 4°C O/N. The beads were then washed with 1X PBS thrice for 10 minutes each on rotation at 4°C. The DNA was then eluted from the protein G beads using DNA elution buffer (1% SDS, 100mM NaHCO3). The DNA was then purified with the help of PCR purification kit (Qiagen). The fold enrichment of MIR503HG was analyzed by qRT-PCR. NOTCH1 and p27 were used as positive controls.

### Proteomic Analysis

The proteomics analysis was performed for MIR503HG knockdown in both the asynchronous and the quiescence conditions. The protein samples for mass spectrometry analysis were prepared using urea lysis buffer. The samples were loaded on Orbitrap LC-MS (Thermo Fisher Scientific). The differential expression data obtained from mass spectrometry was further analyzed with the help of Gene Ontology tools-DAVID and PANTHER.

**Table-1:**
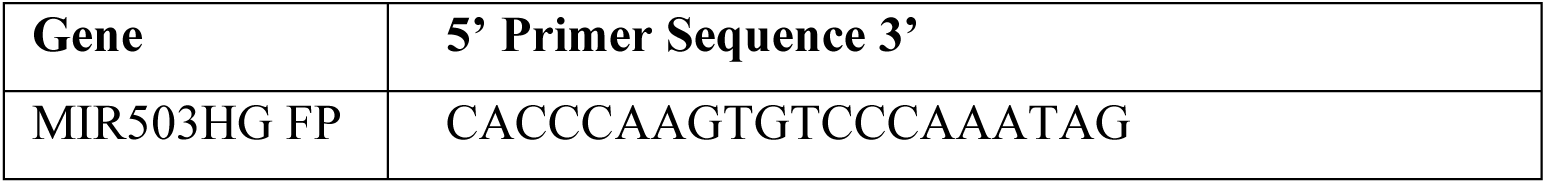
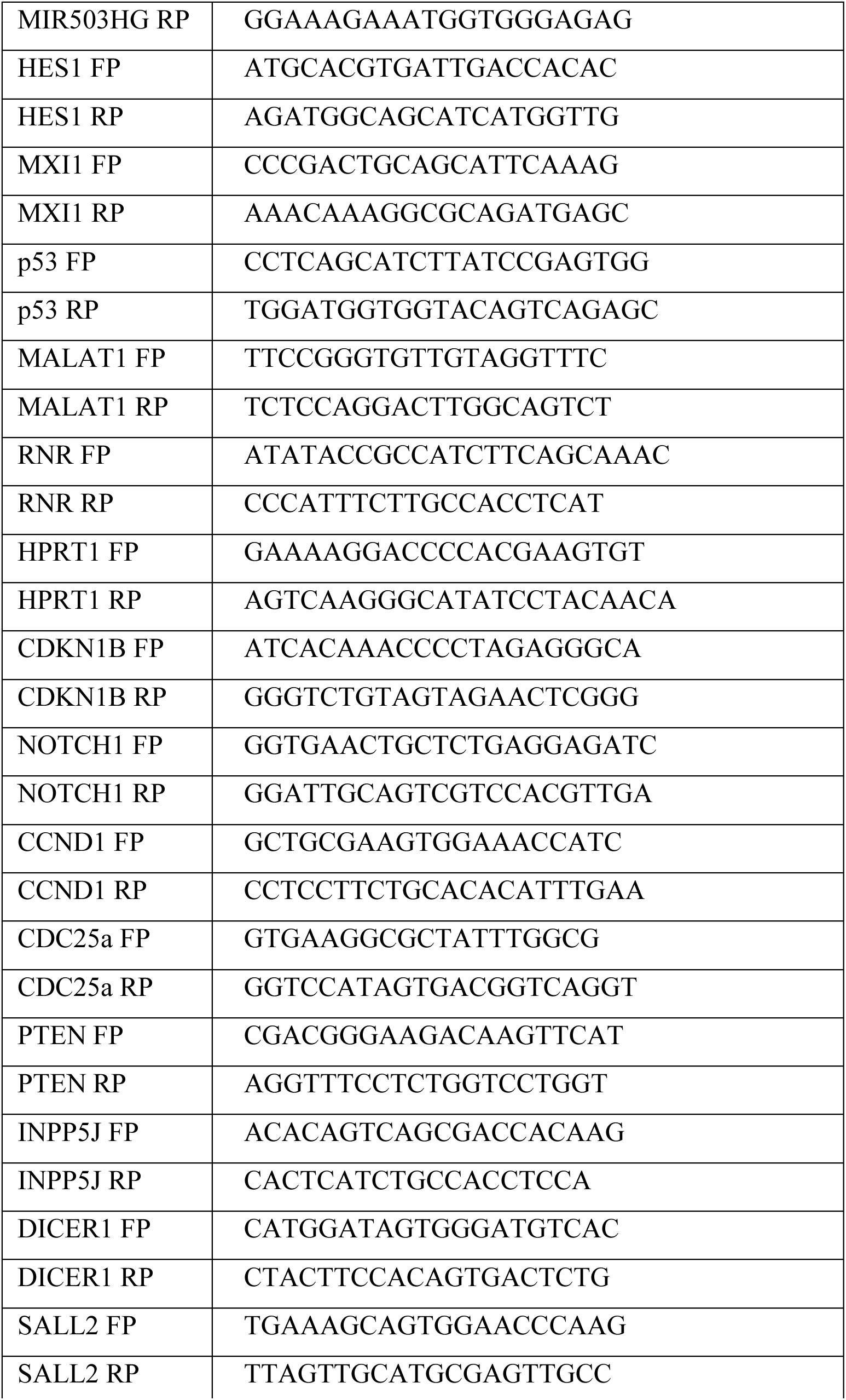
List of primers:

**Table-2:** List of Antibodies.

| Antibody | Dilution | Cat No |
| --- | --- | --- |
| p27 | 1:1000 | sc-1641 |
| Cyclin D1 | 1:1000 | sc-450 |
| CDC25a | 1:500 | sc-7389 |
| PTEN | 1:1000 | sc-7974 |
| pAkt(Thr308) | 1:1000 | 9275S |
| Akt | 1:1000 | 9272 |
| Ago2 | 1:1000 | SAB4200085-25UL |
| α-Tubulin | 1: 50000 | T6199-100UL |
| HRP-linked Caprine Anti-Mouse IgG Polyclonal | 1:10000 | SAA544Mu19 |
| Anti-rabbit IgG, HRP | 1:10000 | 7074 |
| Anti-rat IgG, HRP-linked Antibody | 1:10000 | 7077S |

**Table-3:** miRanda and TargetScan Raw Data:

| miRanda Data Table |  |  |  |  |  |  |  |
| --- | --- | --- | --- | --- | --- | --- | --- |
| miRNA | lncRNA | Aligment score | Energy | Length1 | Length2 | Start | End |
| >>hsa-miR-1343-5p | MIR503HG1 | 176 | -39.06 | 6700 | 22 | 786 | 38 |
| >>hsa-miR-191-5p | MIR503HG1 | 175 | -24.67 | 860 | 23 | 786 | 765 |
| >>hsa-miR-30c-1-3p | MIR503HG1 | 171 | -25.73 | 1072 | 22 | 786 | 452 |
| >>hsa-miR-4765 | MIR503HG1 | 170 | -21.09 | 7276 | 22 | 786 | 378 |
| >>hsa-miR-6777-5p | MIR503HG1 | 166 | -35.06 | 8924 | 23 | 786 | 38 |
| >>hsa-miR-508-3p | MIR503HG1 | 165 | -21.64 | 2120 | 23 | 786 | 278 |
| >>hsa-miR-6868-3p | MIR503HG1 | 165 | -25.04 | 9656 | 21 | 786 | 66 |
| >>hsa-miR-6887-5p | MIR503HG1 | 163 | -35.27 | 9804 | 23 | 786 | 463 |
| >>hsa-miR-5694 | MIR503HG1 | 162 | -25.69 | 8064 | 21 | 786 | 430 |
| >>hsa-miR-939-5p | MIR503HG1 | 161 | -35.43 | 3308 | 24 | 786 | 37 |
| >>hsa-miR-1237-3p | MIR503HG1 | 160 | -26.7 | 3448 | 21 | 786 | 64 |

| TargetScan Data Table |  |  |  |  |  |  |  |
| --- | --- | --- | --- | --- | --- | --- | --- |
| Gene_ID | miRNA family_ID | MSA start | MSA end | UTR start | UTR end | Group num | Site type |
| MIR503HG | miR-1237-3p | 77 | 84 | 77 | 84 | 6447 | 8mer-1a |
| MIR503HG | miR-1287-5p | 183 | 190 | 183 | 190 | 6473 | 8mer-1a |
| MIR503HG | miR-1293/4483 | 351 | 358 | 351 | 358 | 6477 | 8mer-1a |
| MIR503HG | miR-1302/4298 | 361 | 368 | 361 | 368 | 6479 | 8mer-1a |
| MIR503HG | miR-1343-3p | 159 | 166 | 159 | 166 | 6484 | 8mer-1a |
| MIR503HG | miR-142-3p.1 | 234 | 241 | 234 | 241 | 6486 | 8mer-1a |
| MIR503HG | miR-190a-3p | 380 | 387 | 380 | 387 | 6512 | 8mer-1a |
| MIR503HG | miR-191-5p | 779 | 786 | 779 | 786 | 6513 | 8mer-1a |
| MIR503HG | miR-3122/3913-5p | 362 | 369 | 362 | 369 | 6552 | 8mer-1a |
| MIR503HG | miR-3182 | 268 | 275 | 268 | 275 | 6580 | 8mer-1a |
| MIR503HG | miR-4255 | 232 | 239 | 232 | 239 | 6638 | 8mer-1a |
| MIR503HG | miR-4436b-5p | 702 | 709 | 702 | 709 | 6671 | 8mer-1a |
| MIR503HG | miR-4513/6855-3p | 558 | 565 | 558 | 565 | 6691 | 8mer-1a |
| MIR503HG | miR-4725-5p | 634 | 641 | 634 | 641 | 6735 | 8mer-1a |
| MIR503HG | miR-4765 | 391 | 398 | 391 | 398 | 6744 | 8mer-1a |
| MIR503HG | miR-4797-5p | 663 | 670 | 663 | 670 | 6755 | 8mer-1a |
| MIR503HG | miR-489-3p | 622 | 629 | 622 | 629 | 6757 | 8mer-1a |
| MIR503HG | miR-5009-5p/8058 | 453 | 460 | 453 | 460 | 6765 | 8mer-1a |
| MIR503HG | miR-508-3p | 293 | 300 | 293 | 300 | 6775 | 8mer-1a |
| MIR503HG | miR-5584-3p | 694 | 701 | 694 | 701 | 6817 | 8mer-1a |
| MIR503HG | miR-5689 | 675 | 682 | 675 | 682 | 6821 | 8mer-1a |
| MIR503HG | miR-5692bc | 404 | 411 | 404 | 411 | 6823 | 8mer-1a |
| MIR503HG | miR-625-5p | 495 | 502 | 495 | 502 | 6855 | 8mer-1a |
| MIR503HG | miR-653-5p | 280 | 287 | 280 | 287 | 6874 | 8mer-1a |
| MIR503HG | miR-6731-5p/8085 | 467 | 474 | 467 | 474 | 6881 | 8mer-1a |
| MIR503HG | miR-6742-3p | 158 | 165 | 158 | 165 | 6887 | 8mer-1a |
| MIR503HG | miR-6742-5p | 709 | 716 | 709 | 716 | 6888 | 8mer-1a |
| MIR503HG | miR-6795-5p/6887-5p | 218 | 225 | 218 | 225 | 6907 | 8mer-1a |
| MIR503HG | miR-6808-3p | 656 | 663 | 656 | 663 | 6914 | 8mer-1a |
| MIR503HG | miR-6828-3p | 74 | 81 | 74 | 81 | 6920 | 8mer-1a |
| MIR503HG | miR-6856-3p | 729 | 736 | 729 | 736 | 6929 | 8mer-1a |
| MIR503HG | miR-6868-3p | 78 | 85 | 78 | 85 | 6932 | 8mer-1a |
| MIR503HG | miR-7109-5p | 219 | 226 | 219 | 226 | 6941 | 8mer-1a |
| MIR503HG | miR-7152-5p | 716 | 723 | 716 | 723 | 6946 | 8mer-1a |
| MIR503HG | miR-766-5p | 135 | 142 | 135 | 142 | 6955 | 8mer-1a |
| MIR503HG | miR-7973 | 636 | 643 | 636 | 643 | 6960 | 8mer-1a |
| MIR503HG | miR-1207-5p/4763-3p | 35 | 41 | 35 | 41 | 6438 | 7mer-1a |
| MIR503HG | miR-1231 | 689 | 695 | 689 | 695 | 6442 | 7mer-1a |
| MIR503HG | miR-1236-5p | 392 | 398 | 392 | 398 | 6445 | 7mer-1a |
| MIR503HG | miR-1248 | 78 | 84 | 78 | 84 | 6455 | 7mer-1a |
| MIR503HG | miR-1262/4701-3p/6736-5p | 353 | 359 | 353 | 359 | 6459 | 7mer-1a |
| MIR503HG | miR-1269 | 454 | 460 | 454 | 460 | 6461 | 7mer-1a |
| MIR503HG | miR-1273g-3p | 141 | 147 | 141 | 147 | 6465 | 7mer-1a |
| MIR503HG | miR-1275/4665-5p | 220 | 226 | 220 | 226 | 6469 | 7mer-1a |
| MIR503HG | miR-1305 | 345 | 351 | 345 | 351 | 6481 | 7mer-1a |
| MIR503HG | miR-138-2-3p | 367 | 373 | 367 | 373 | 6485 | 7mer-1a |
| MIR503HG | miR-148b-5p/6874-3p | 695 | 701 | 695 | 701 | 6493 | 7mer-1a |
| MIR503HG | miR-181a-2-3p | 141 | 147 | 141 | 147 | 6503 | 7mer-1a |
| MIR503HG | miR-1908-5p | 546 | 552 | 546 | 552 | 6511 | 7mer-1a |
| MIR503HG | miR-1914-5p | 715 | 721 | 715 | 721 | 6515 | 7mer-1a |
| MIR503HG | miR-193b-5p | 708 | 714 | 708 | 714 | 6517 | 7mer-1a |
| MIR503HG | miR-198 | 262 | 268 | 262 | 268 | 6520 | 7mer-1a |
| MIR503HG | miR-212-5p | 457 | 463 | 457 | 463 | 6526 | 7mer-1a |
| MIR503HG | miR-219-5p | 294 | 300 | 294 | 300 | 6527 | 7mer-1a |
| MIR503HG | miR-2681-5p | 668 | 674 | 668 | 674 | 6535 | 7mer-1a |
| MIR503HG | miR-26b-3p | 588 | 594 | 588 | 594 | 6536 | 7mer-1a |
| MIR503HG | miR-3074-5p | 717 | 723 | 717 | 723 | 6545 | 7mer-1a |
| MIR503HG | miR-3121-5p | 227 | 233 | 227 | 233 | 6550 | 7mer-1a |
| MIR503HG | miR-3135a | 338 | 344 | 338 | 344 | 6558 | 7mer-1a |
| MIR503HG | miR-3135b | 184 | 190 | 184 | 190 | 6559 | 7mer-1a |
| MIR503HG | miR-3147 | 222 | 228 | 222 | 228 | 6563 | 7mer-1a |
| MIR503HG | miR-3151-5p | 710 | 716 | 710 | 716 | 6567 | 7mer-1a |
| MIR503HG | miR-3168 | 695 | 701 | 695 | 701 | 6575 | 7mer-1a |
| MIR503HG | miR-3171 | 238 | 244 | 238 | 244 | 6577 | 7mer-1a |
| MIR503HG | miR-3189-3p | 457 | 463 | 457 | 463 | 6584 | 7mer-1a |
| MIR503HG | miR-328-5p/6885-5p | 219 | 225 | 219 | 225 | 6590 | 7mer-1a |
| MIR503HG | miR-363-5p/6745 | 352 | 358 | 352 | 358 | 6604 | 7mer-1a |
| MIR503HG | miR-3652/4430 | 94 | 100 | 94 | 100 | 6606 | 7mer-1a |
| MIR503HG | miR-3655 | 691 | 697 | 691 | 697 | 6612 | 7mer-1a |
| MIR503HG | miR-3657 | 264 | 270 | 264 | 270 | 6613 | 7mer-1a |
| MIR503HG | miR-3662 | 326 | 332 | 326 | 332 | 6614 | 7mer-1a |
| MIR503HG | miR-3671 | 584 | 590 | 584 | 590 | 6616 | 7mer-1a |
| MIR503HG | miR-3675-5p | 498 | 504 | 498 | 504 | 6617 | 7mer-1a |
| MIR503HG | miR-369-3p | 405 | 411 | 405 | 411 | 6620 | 7mer-1a |
| MIR503HG | miR-3691-3p | 609 | 615 | 609 | 615 | 6621 | 7mer-1a |
| MIR503HG | miR-377-5p/6086 | 573 | 579 | 573 | 579 | 6624 | 7mer-1a |
| MIR503HG | miR-3911 | 712 | 718 | 712 | 718 | 6630 | 7mer-1a |
| MIR503HG | miR-3913-3p | 622 | 628 | 622 | 628 | 6632 | 7mer-1a |
| MIR503HG | miR-4268 | 72 | 78 | 72 | 78 | 6644 | 7mer-1a |
| MIR503HG | miR-4297/5581-5p | 177 | 183 | 177 | 183 | 6652 | 7mer-1a |
| MIR503HG | miR-4324 | 69 | 75 | 69 | 75 | 6661 | 7mer-1a |
| MIR503HG | miR-4326 | 230 | 236 | 230 | 236 | 6662 | 7mer-1a |
| MIR503HG | miR-4425 | 222 | 228 | 222 | 228 | 6665 | 7mer-1a |
| MIR503HG | miR-4433a-5p | 746 | 752 | 746 | 752 | 6667 | 7mer-1a |
| MIR503HG | miR-4445-5p | 294 | 300 | 294 | 300 | 6673 | 7mer-1a |
| MIR503HG | miR-4447/4472 | 710 | 716 | 710 | 716 | 6675 | 7mer-1a |
| MIR503HG | miR-4448 | 72 | 78 | 72 | 78 | 6676 | 7mer-1a |
| MIR503HG | miR-4505/5787 | 94 | 100 | 94 | 100 | 6687 | 7mer-1a |
| MIR503HG | miR-450a-1-3p | 363 | 369 | 363 | 369 | 6689 | 7mer-1a |
| MIR503HG | miR-4537 | 66 | 72 | 66 | 72 | 6698 | 7mer-1a |
| MIR503HG | miR-4634 | 122 | 128 | 122 | 128 | 6699 | 7mer-1a |
| MIR503HG | miR-4635 | 283 | 289 | 283 | 289 | 6700 | 7mer-1a |
| MIR503HG | miR-4652-3p | 641 | 647 | 641 | 647 | 6705 | 7mer-1a |
| MIR503HG | miR-4666a-5p | 529 | 535 | 529 | 535 | 6709 | 7mer-1a |
| MIR503HG | miR-4669 | 264 | 270 | 264 | 270 | 6715 | 7mer-1a |
| MIR503HG | miR-4677-3p | 639 | 645 | 639 | 645 | 6716 | 7mer-1a |
| MIR503HG | miR-4679 | 639 | 645 | 639 | 645 | 6717 | 7mer-1a |
| MIR503HG | miR-4683 | 190 | 196 | 190 | 196 | 6719 | 7mer-1a |
| MIR503HG | miR-4723-5p/5698/6870-5p/7111-5p | 496 | 502 | 496 | 502 | 6734 | 7mer-1a |
| MIR503HG | miR-4730 | 547 | 553 | 547 | 553 | 6737 | 7mer-1a |
| MIR503HG | miR-4736 | 35 | 41 | 35 | 41 | 6738 | 7mer-1a |
| MIR503HG | miR-4743-3p | 641 | 647 | 641 | 647 | 6740 | 7mer-1a |
| MIR503HG | miR-4762-5p | 583 | 589 | 583 | 589 | 6742 | 7mer-1a |
| MIR503HG | miR-491-5p | 710 | 716 | 710 | 716 | 6758 | 7mer-1a |
| MIR503HG | miR-494-3p | 280 | 286 | 280 | 286 | 6759 | 7mer-1a |
| MIR503HG | miR-5002-5p | 365 | 371 | 365 | 371 | 6763 | 7mer-1a |
| MIR503HG | miR-5010-5p | 496 | 502 | 496 | 502 | 6768 | 7mer-1a |
| MIR503HG | miR-5011-5p | 380 | 386 | 380 | 386 | 6770 | 7mer-1a |
| MIR503HG | miR-502-5p | 458 | 464 | 458 | 464 | 6771 | 7mer-1a |
| MIR503HG | miR-503-3p | 502 | 508 | 502 | 508 | 6772 | 7mer-1a |
| MIR503HG | miR-526b-5p | 284 | 290 | 284 | 290 | 6790 | 7mer-1a |
| MIR503HG | miR-543 | 423 | 429 | 423 | 429 | 6796 | 7mer-1a |
| MIR503HG | miR-543 | 603 | 609 | 603 | 609 | 6798 | 7mer-1a |
| MIR503HG | miR-544b | 69 | 75 | 69 | 75 | 6800 | 7mer-1a |
| MIR503HG | miR-5683 | 447 | 453 | 447 | 453 | 6820 | 7mer-1a |
| MIR503HG | miR-578 | 285 | 291 | 285 | 291 | 6830 | 7mer-1a |
| MIR503HG | miR-582-3p | 194 | 200 | 194 | 200 | 6833 | 7mer-1a |
| MIR503HG | miR-593-5p | 90 | 96 | 90 | 96 | 6839 | 7mer-1a |
| MIR503HG | miR-6071 | 75 | 81 | 75 | 81 | 6845 | 7mer-1a |
| MIR503HG | miR-6073 | 235 | 241 | 235 | 241 | 6846 | 7mer-1a |
| MIR503HG | miR-632/4288 | 689 | 695 | 689 | 695 | 6856 | 7mer-1a |
| MIR503HG | miR-642-3p | 597 | 603 | 597 | 603 | 6858 | 7mer-1a |
| MIR503HG | miR-644a | 712 | 718 | 712 | 718 | 6861 | 7mer-1a |
| MIR503HG | miR-6504-3p | 400 | 406 | 400 | 406 | 6867 | 7mer-1a |
| MIR503HG | miR-6509-3p | 703 | 709 | 703 | 709 | 6869 | 7mer-1a |
| MIR503HG | miR-6513-5p | 363 | 369 | 363 | 369 | 6871 | 7mer-1a |
| MIR503HG | miR-654-5p | 471 | 477 | 471 | 477 | 6875 | 7mer-1a |
| MIR503HG | miR-655-5p | 573 | 579 | 573 | 579 | 6876 | 7mer-1a |
| MIR503HG | miR-656-3p | 378 | 384 | 378 | 384 | 6877 | 7mer-1a |
| MIR503HG | miR-6741-5p | 353 | 359 | 353 | 359 | 6886 | 7mer-1a |
| MIR503HG | miR-6748-5p | 504 | 510 | 504 | 510 | 6889 | 7mer-1a |
| MIR503HG | miR-6751-5p/6803-5p | 220 | 226 | 220 | 226 | 6890 | 7mer-1a |
| MIR503HG | miR-6756-5p/6766-5p | 352 | 358 | 352 | 358 | 6892 | 7mer-1a |
| MIR503HG | miR-6757-3p | 397 | 403 | 397 | 403 | 6893 | 7mer-1a |
| MIR503HG | miR-6769a-5p/6769b-5p | 471 | 477 | 471 | 477 | 6901 | 7mer-1a |
| MIR503HG | miR-6770-3p | 547 | 553 | 547 | 553 | 6902 | 7mer-1a |
| MIR503HG | miR-6793-5p | 504 | 510 | 504 | 510 | 6906 | 7mer-1a |
| MIR503HG | miR-6843-3p/6848-3p | 112 | 118 | 112 | 118 | 6925 | 7mer-1a |
| MIR503HG | miR-6852-5p | 159 | 165 | 159 | 165 | 6927 | 7mer-1a |
| MIR503HG | miR-6856-3p | 310 | 316 | 310 | 316 | 6928 | 7mer-1a |
| MIR503HG | miR-6857-3p | 559 | 565 | 559 | 565 | 6930 | 7mer-1a |
| MIR503HG | miR-6878-5p | 468 | 474 | 468 | 474 | 6934 | 7mer-1a |
| MIR503HG | miR-6890-5p | 498 | 504 | 498 | 504 | 6937 | 7mer-1a |
| MIR503HG | miR-6894-5p | 136 | 142 | 136 | 142 | 6938 | 7mer-1a |
| MIR503HG | miR-7154-3p | 136 | 142 | 136 | 142 | 6947 | 7mer-1a |
| MIR503HG | miR-765 | 136 | 142 | 136 | 142 | 6954 | 7mer-1a |
| MIR503HG | miR-767-3p | 74 | 80 | 74 | 80 | 6956 | 7mer-1a |
| MIR503HG | miR-7848-3p | 374 | 380 | 374 | 380 | 6957 | 7mer-1a |
| MIR503HG | miR-7973 | 656 | 662 | 656 | 662 | 6961 | 7mer-1a |
| MIR503HG | miR-873-5p.1 | 104 | 110 | 104 | 110 | 6967 | 7mer-1a |
| MIR503HG | miR-887-5p | 363 | 369 | 363 | 369 | 6971 | 7mer-1a |
| MIR503HG | miR-92a-2-5p | 471 | 477 | 471 | 477 | 6978 | 7mer-1a |
| MIR503HG | miR-939-3p | 159 | 165 | 159 | 165 | 6979 | 7mer-1a |

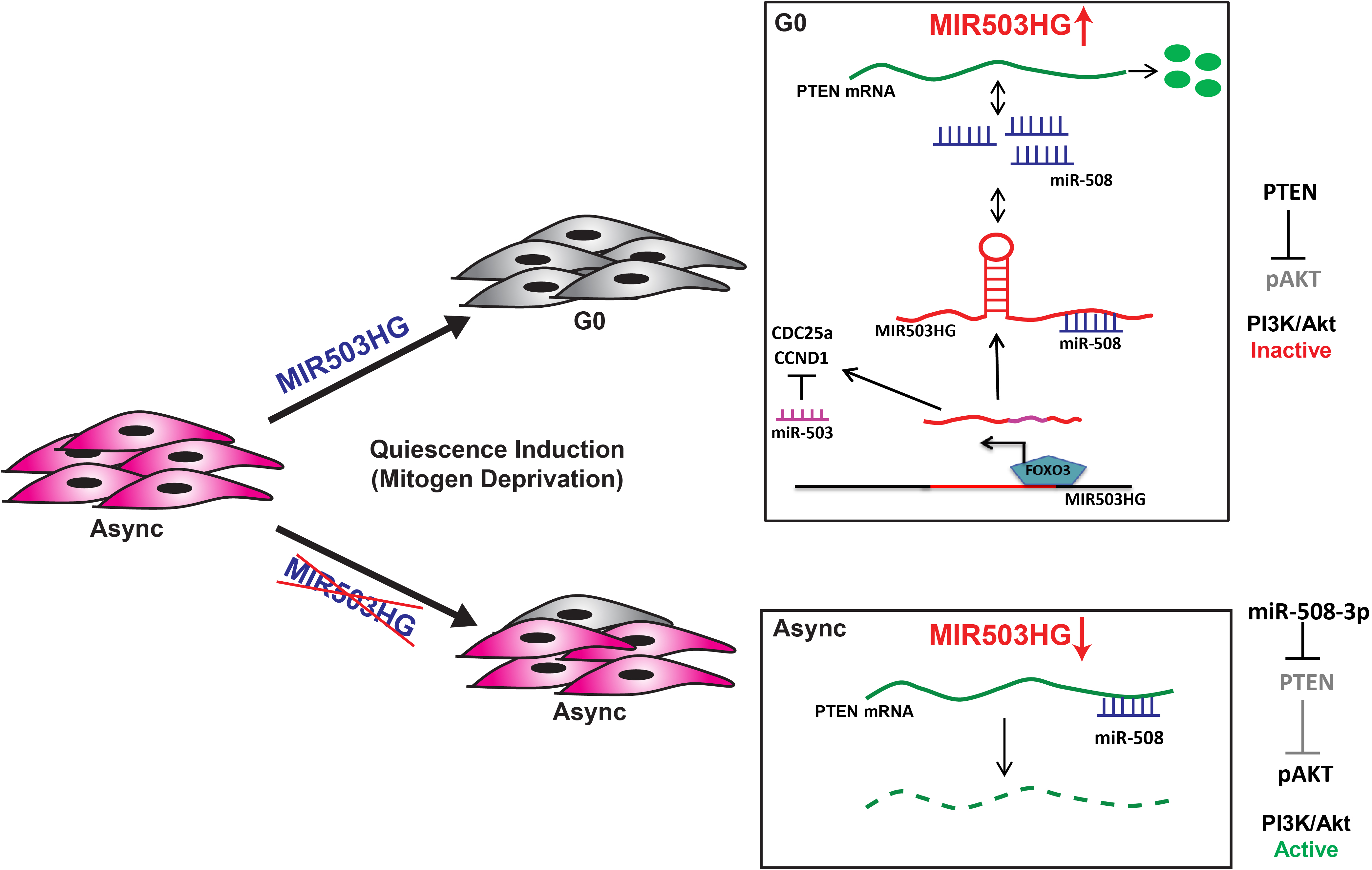

**Fig. S1.**
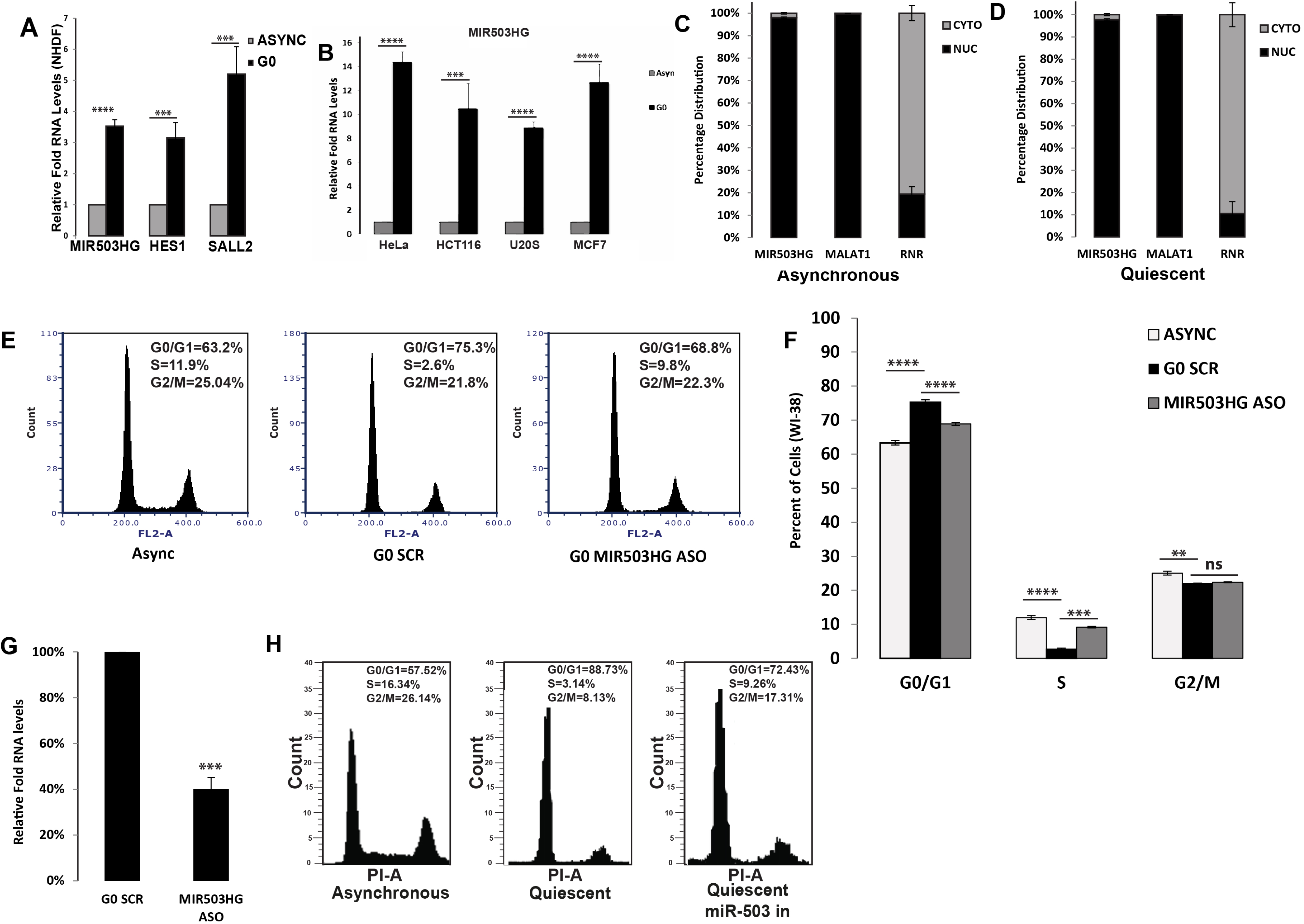
(A) qRT-PCR analysis of MIR503HG, HES1 and SALL2 RNA in asynchronous and G0 NHDFs (Normal human dermal fibroblasts). (B) qRT-PCR analysis of MIR503HG in asynchronous and G0 Hela, HCT116, U2OS and MCF7 cells. (C, D) Subcellular fractionation followed by qRT-PCR analysis for MIR503HG, MALAT1 and RNR in asynchronous and quiescent WI38 cells. (E) PI-Flow cytometry analysis of asynchronous and quiescent WI38 cells treated with scr-oligo and MIR503HG-ASO. (F) Quantitative representation of percentage cell population from PI-Flow cytometry analysis of asynchronous and quiescent WI38 cells treated with scr-oligo and MIR503HG-ASO. (G) qRT-PCR analysis for miR503 in G0 control and MIR503HG depleted cells. (H) PI-Flow cytometry analysis of asynchronous, quiescent and WI38 cells treated with miR503 inhibitor.

**Fig. S2.**
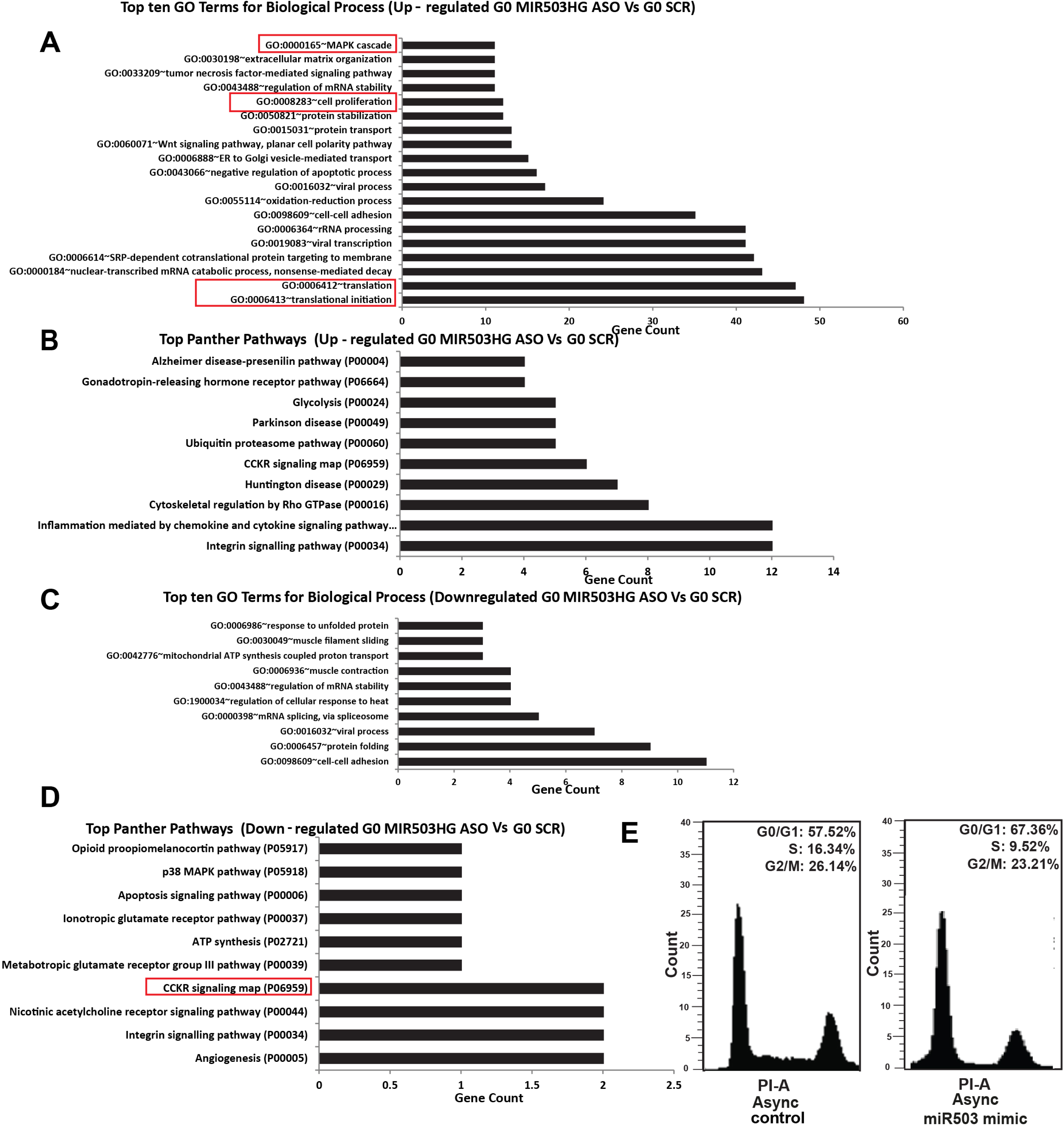
Top significant biological processes for genes whose protein levels are upregulated in MIR503HG depleted WI38 cells at G0. (B) Top significant biological processes for genes whose protein levels are downregulated in MIR503HG depleted WI38 cells at G0. (C) Top significant panther pathways upregulated in MIR503HG depleted WI38 cells at G0. (D) Top significant panther pathways downregulated in MIR503HG depleted WI38 cells at G0. (E) PI-Flow cytometry analysis of asynchronous control and miR503 mimic treated WI38 cells.

**Fig. S3.**
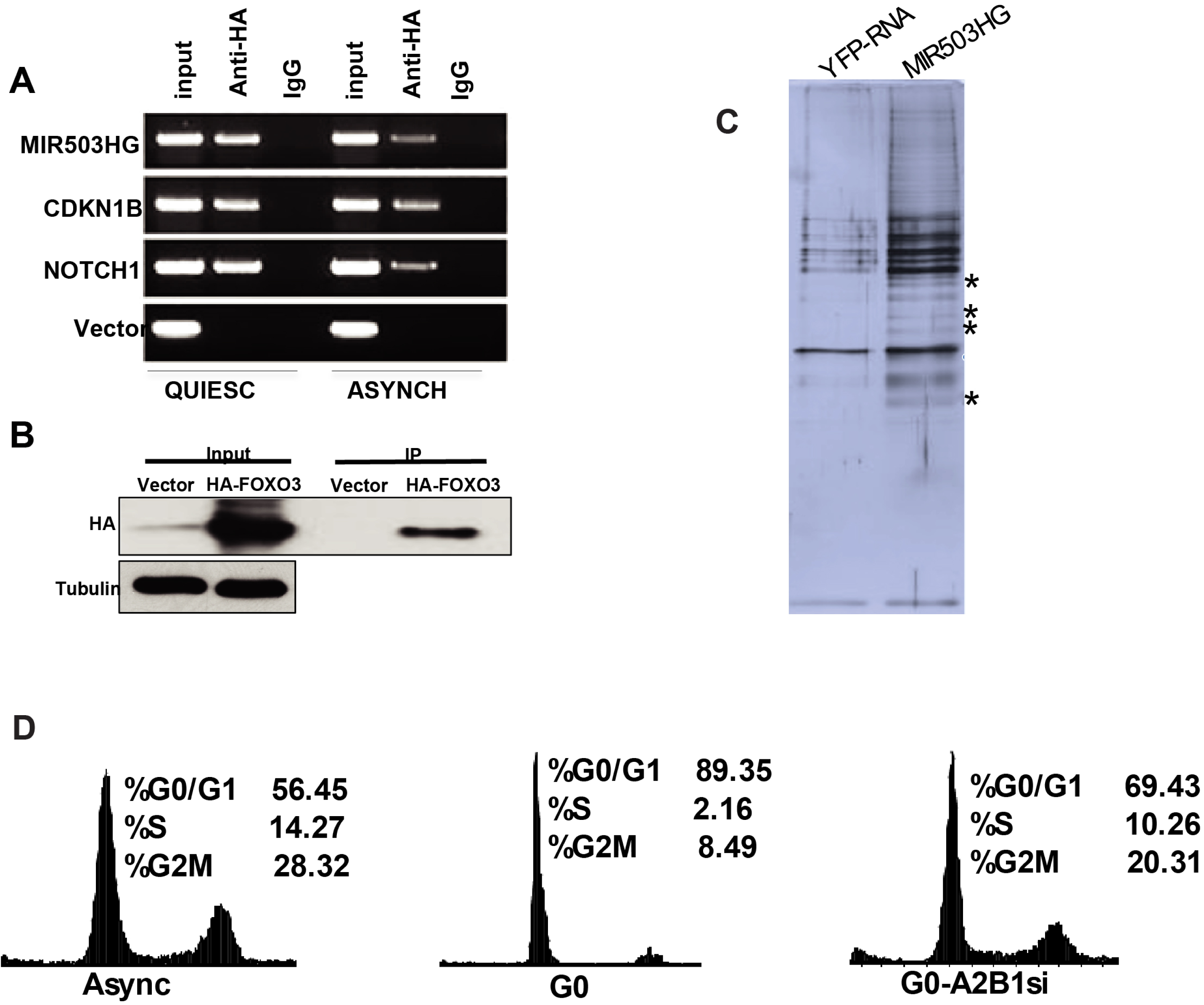
(A) Semi-quantitative PCR to analyze relative fold enrichment of FOXO3 at MIR503HG, CDKN1B and NOTCH1 promoters in asynchronous and G0 cells through chromatin immunoprecipitation using HA antibody. (B) Western blot analysis of ChIP assay using HA antibody in control and HA-FOXO3 transfected cells. (C) Gel picture demonstrating the in vitro transcribed MIR503HG RNA. (D) Silver-stained gel image of RNA pull down assay with YFP-RNA and MIR503HG. (E) PI-Flow cytometry analysis of asynchronously growing, G0 and G0 cells treated with hnRNPA2B1-siRNA WI38 cells.

**Fig. S4.**
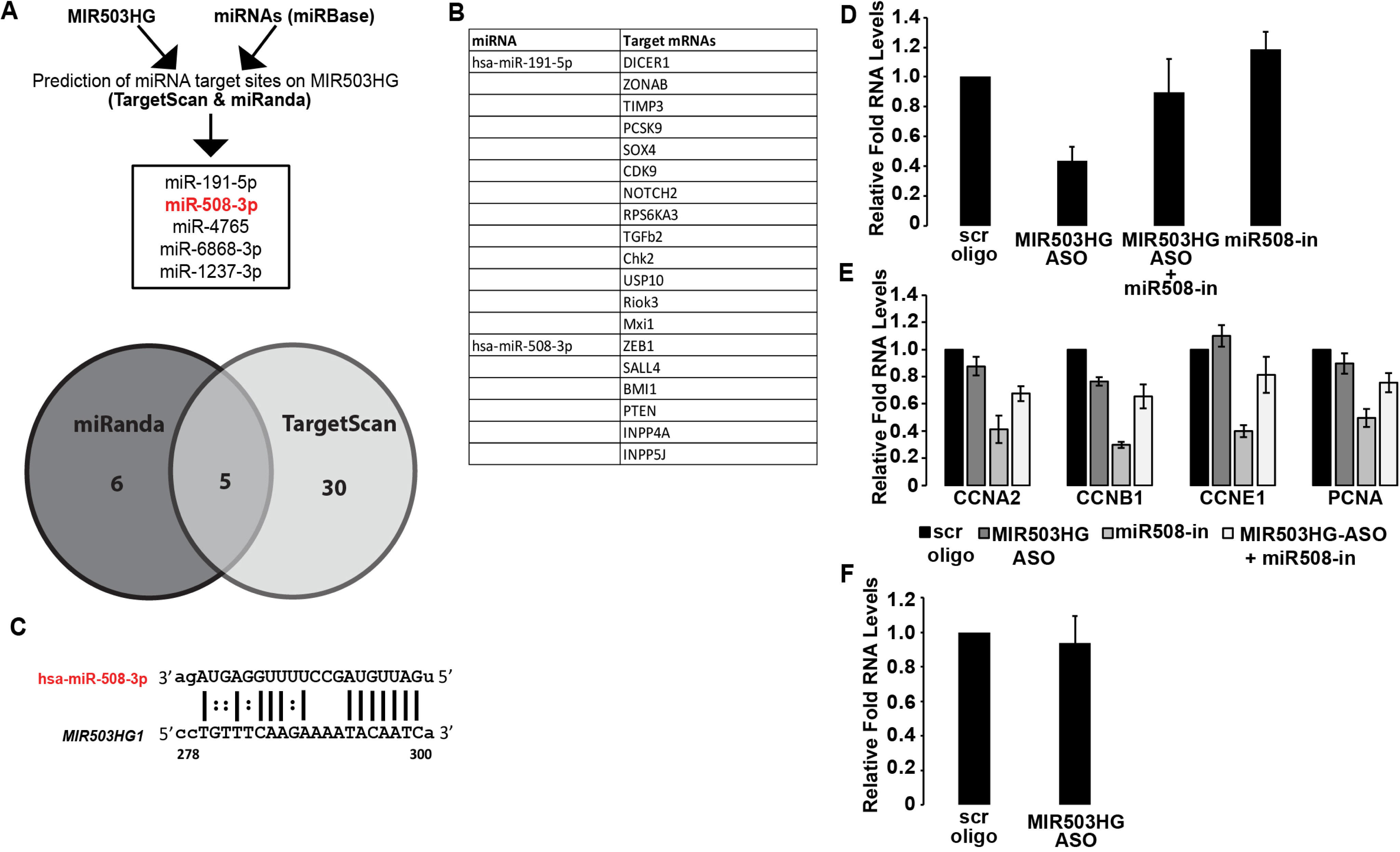
(A) Schematic showing the strategy employed to predict putative binding sites for miRNAs on MIR503HG transcript. (B) List of known mRNA targets for miR191 and miR508. (C) Complementarity based binding of miR508 with MIR503HG RNA.

## Notes

### Competing Interest Statement

The authors have declared no competing interest.

