## Supplementary material for "FOXO3 regulated MIR503HG safeguards cellular quiescence by modulating PI3K/Akt pathway via miR-508/PTEN axis"

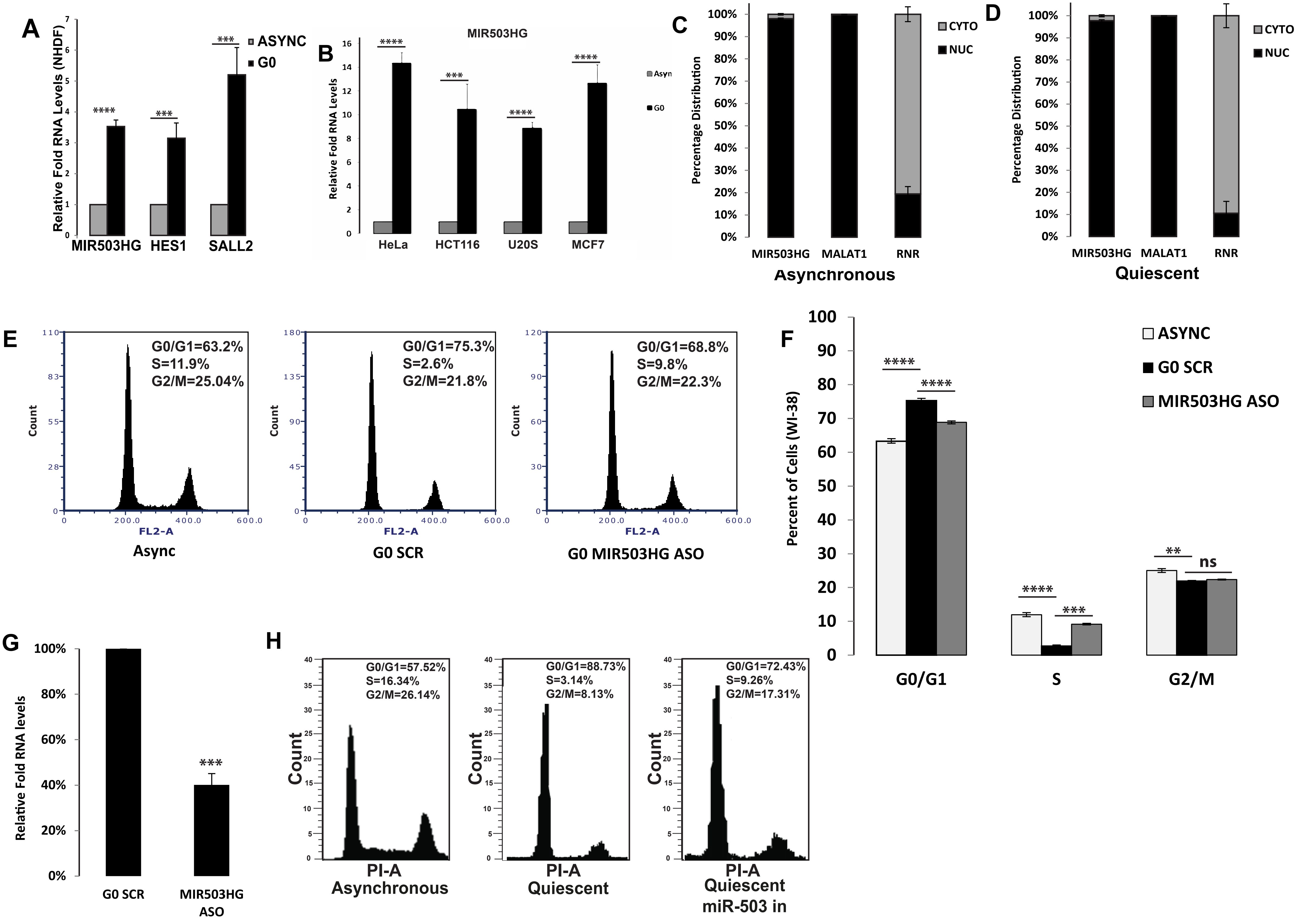
A
Relative Fold RNA Levels (NHDF)
ASYNC
G0
MIR503HG
HES1
SALL2
B
MIR503HG
Relative Fold RNA Levels
Asyn
G0
HeLa
HCT116
U20S
MCF7
C
Percentage Distribution
CYTO
NUC
MIR503HG
MALAT1
RNR
Asynchronous
D
Percentage Distribution
CYTO
NUC
MIR503HG
MALAT1
RNR
Quiescent
E
G0/G1=63.2%
S=11.9%
G2/M=25.04%
Count
FL2-A
Async
G0/G1=75.3%
S=2.6%
G2/M=21.8%
Count
FL2-A
G0 SCR
G0/G1=68.8%
S=9.8%
G2/M=22.3%
Count
FL2-A
G0 MIR503HG ASO
F
Percent of Cells (WI-38)
ASYNC
G0 SCR
MIR503HG ASO
ns
G0/G1
S
G2/M
G
Relative Fold RNA levels
G0 SCR
MIR503HG ASO
H
G0/G1=57.52%
S=16.34%
G2/M=26.14%
Count
PI-A
Asynchronous
G0/G1=88.73%
S=3.14%
G2/M=8.13%
Count
PI-A
Quiescent
G0/G1=72.43%
S=9.26%
G2/M=17.31%
Count
PI-A
Quiescent miR-503 in

Suppl-Fig1

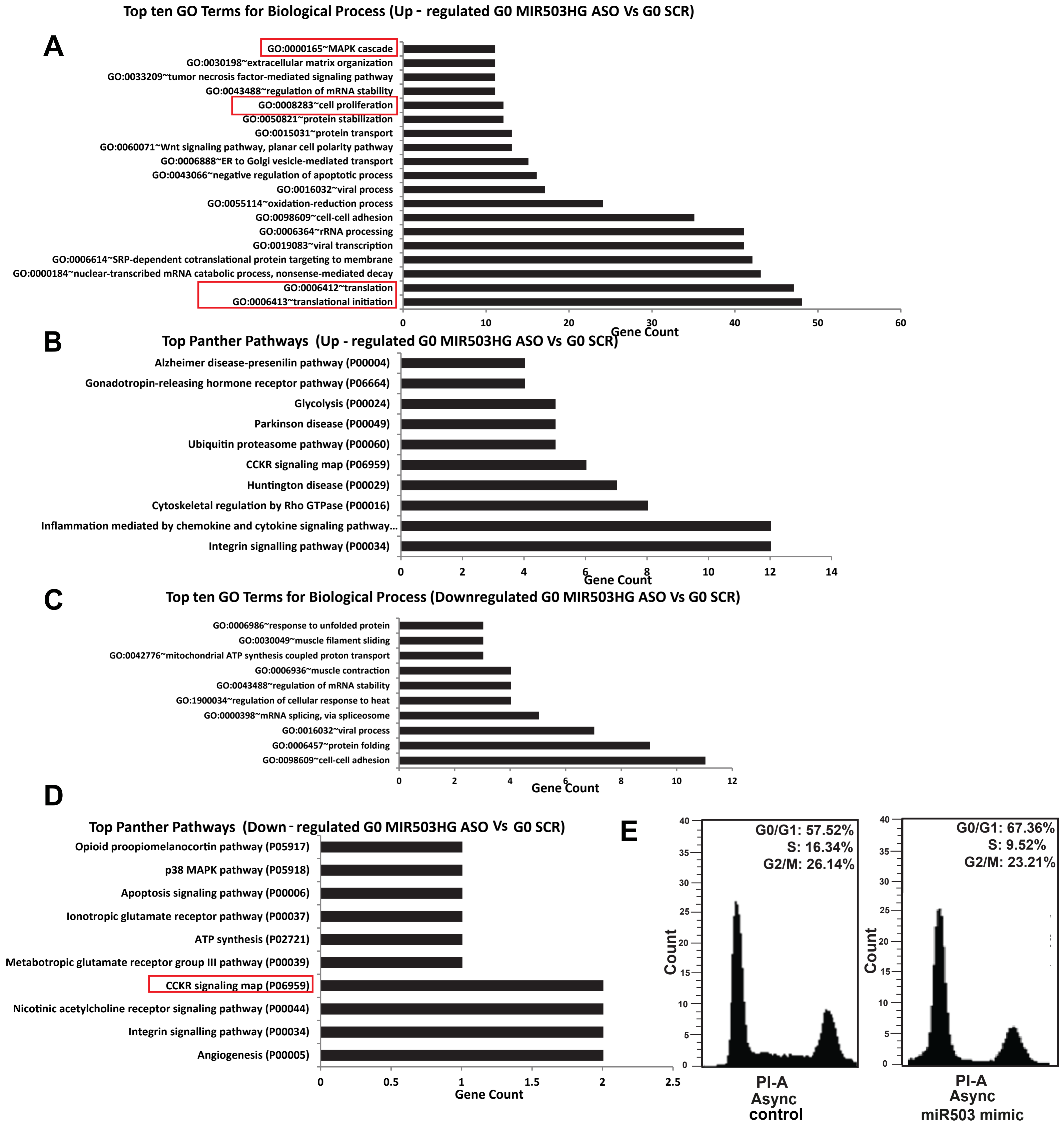
Top ten GO Terms for Biological Process (Up - regulated G0 MIR503HG ASO Vs G0 SCR)
A
GO:0000165~MAPK cascade
GO:0030198~extracellular matrix organization
GO:0033209~tumor necrosis factor-mediated signaling pathway
GO:0043488~regulation of mRNA stability
GO:0008283~cell proliferation
GO:0050821~protein stabilization
GO:0015031~protein transport
GO:0060071~Wnt signaling pathway, planar cell polarity pathway
GO:0006888~ER to Golgi vesicle-mediated transport
GO:0043066~negative regulation of apoptotic process
GO:0016032~viral process
GO:0055114~oxidation-reduction process
GO:0098609~cell-cell adhesion
GO:0006364~rRNA processing
GO:0019083~viral transcription
GO:0006614~SRP-dependent cotranslational protein targeting to membrane
GO:0000184~nuclear-transcribed mRNA catabolic process, nonsense-mediated decay
GO:0006412~translation
GO:0006413~translational initiation
0 10 20 30 40 50 60
Gene Count
B
Top Panther Pathways (Up - regulated G0 MIR503HG ASO Vs G0 SCR)
Alzheimer disease-presenilin pathway (P00004)
Gonadotropin-releasing hormone receptor pathway (P06664)
Glycolysis (P00024)
Parkinson disease (P00049)
Ubiquitin proteasome pathway (P00060)
CCKR signaling map (P06959)
Huntington disease (P00029)
Cytoskeletal regulation by Rho GTPase (P00016)
Inflammation mediated by chemokine and cytokine signaling pathway...
Integrin signalling pathway (P00034)
0 2 4 6 8 10 12 14
Gene Count
C
Top ten GO Terms for Biological Process (Downregulated G0 MIR503HG ASO Vs G0 SCR)
GO:0006986~response to unfolded protein
GO:0030049~muscle filament sliding
GO:0042776~mitochondrial ATP synthesis coupled proton transport
GO:0006936~muscle contraction
GO:0043488~regulation of mRNA stability
GO:1900034~regulation of cellular response to heat
GO:0000398~mRNA splicing, via spliceosome
GO:0016032~viral process
GO:0006457~protein folding
GO:0098609~cell-cell adhesion
0 2 4 6 8 10 12
Gene Count
D
Top Panther Pathways (Down - regulated G0 MIR503HG ASO Vs G0 SCR)
Opioid proopiomelanocortin pathway (P05917)
p38 MAPK pathway (P05918)
Apoptosis signaling pathway (P00006)
Ionotropic glutamate receptor pathway (P00037)
ATP synthesis (P02721)
Metabotropic glutamate receptor group III pathway (P00039)
CCKR signaling map (P06959)
Nicotinic acetylcholine receptor signaling pathway (P00044)
Integrin signalling pathway (P00034)
Angiogenesis (P00005)
0 0.5 1 1.5 2 2.5
Gene Count
E
G0/G1: 57.52%
S: 16.34%
G2/M: 26.14%
Count
PI-A
Async
control
G0/G1: 67.36%
S: 9.52%
G2/M: 23.21%
Count
PI-A
Async
miR503 mimic

Suppl-Fig2

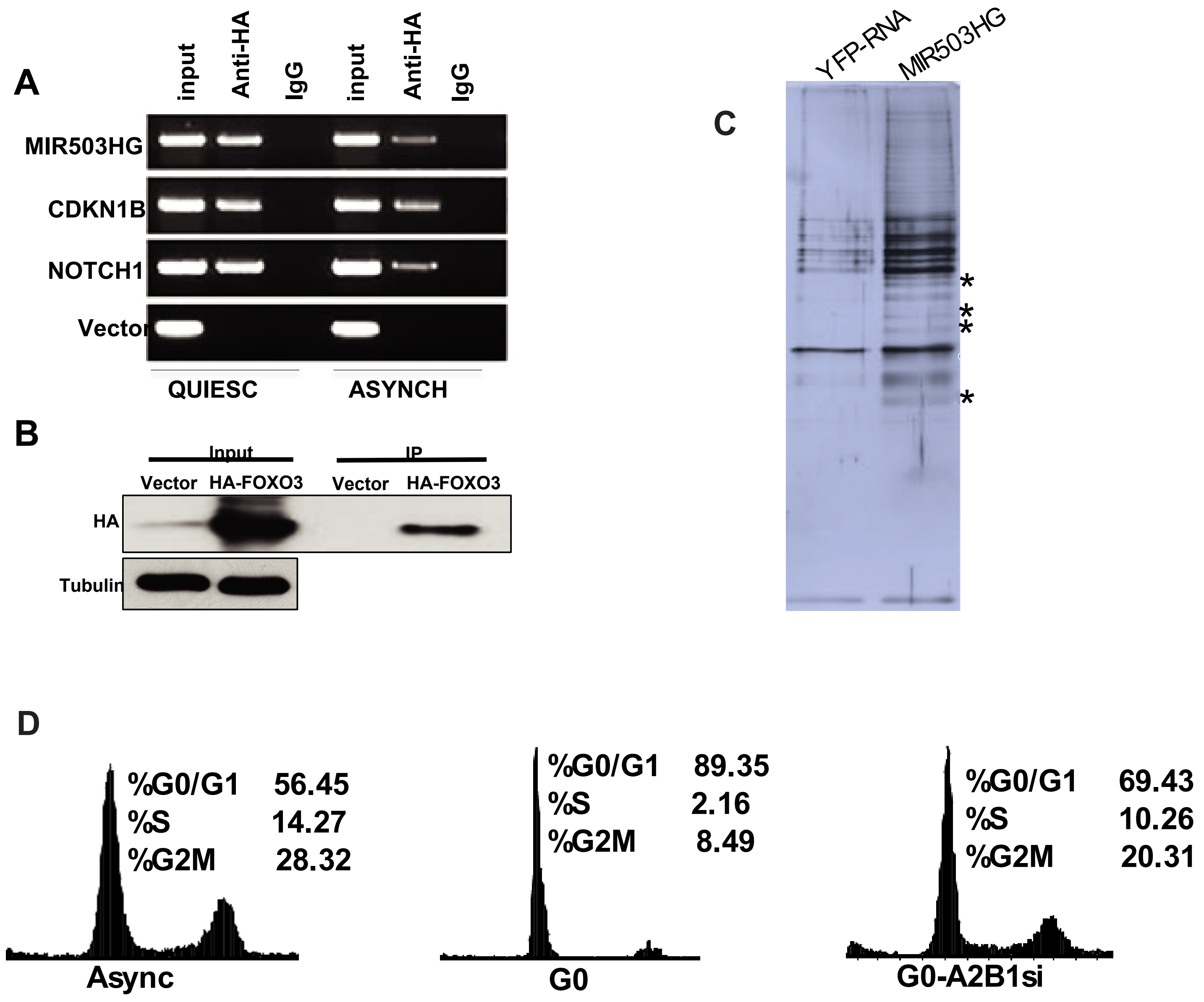
A
input
Anti-HA
IgG
input
Anti-HA
IgG
MIR503HG
CDKN1B
NOTCH1
Vector
QUIESC
ASYNCH
B
Input
IP
Vector
HA-FOXO3
Vector
HA-FOXO3
HA
Tubulin
C
YFP-RNA
MIR503HG
D
%G0/G1 56.45
%S 14.27
%G2M 28.32
Async
%G0/G1 89.35
%S 2.16
%G2M 8.49
G0
%G0/G1 69.43
%S 10.26
%G2M 20.31
G0-A2B1si

Suppl-Fig3

A

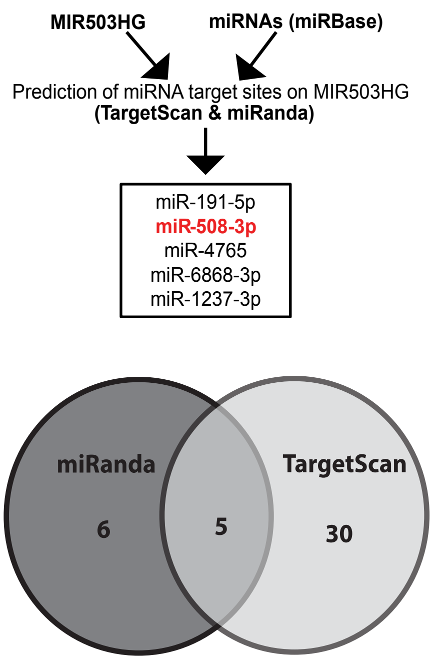


B

| miRNA | Target mRNAs |
|---|---|
| hsa-miR-191-5p | DICER1 |
| | ZONAB |
| | TIMP3 |
| | PCSK9 |
| | SOX4 |
| | CDK9 |
| | NOTCH2 |
| | RPS6KA3 |
| | TGFb2 |
| | Chk2 |
| | USP10 |
| | Riok3 |
| | Mxi1 |
| hsa-miR-508-3p | ZEB1 |
| | SALL4 |
| | BMI1 |
| | PTEN |
| | INPP4A |
| | INPP5J |

C

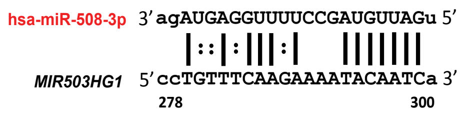


D
E
F

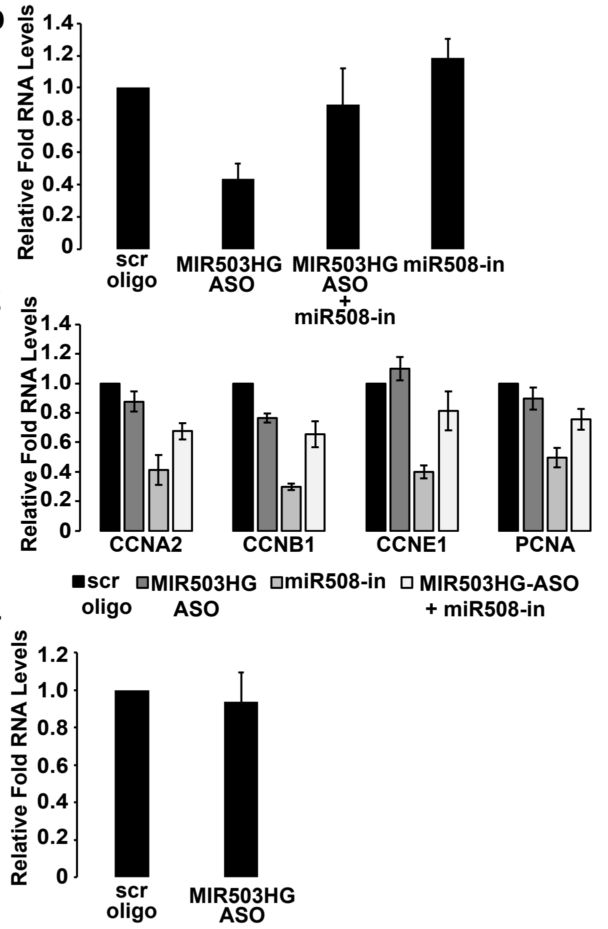


Suppl-Fig4
